# stem.MAP: Identification of cell-type-specific protein complexes in mammalian embryonic stem cells

**DOI:** 10.64898/2026.11.04.764359

**Authors:** Akshaya Rajaraman, Samantha N. Fischer, Miles D. Woodcock-Girard, Riddhiman K. Garge, Kevin Drew

## Abstract

Protein complexes carry out critical functions during development. Mapping cell-type-specific protein complexes in early embryogenesis is essential for uncovering the molecular mechanisms of embryonic development. We unfortunately currently lack a catalog of protein complexes specific for embryogenesis including embryonic stem cells. Here we develop a proteomics-based machine learning workflow to predict protein complexes present in mouse embryonic stem cells (mESC). Our method integrates data from a cell-type agnostic protein complex resource with co-fractionation mass spectrometry (CF-MS) experiments on mESC. Our resource, stem.MAP, identifies 3,986 complexes encompassing ∼133,000 unique interactions, and 8,616 proteins. stem.MAP captures variant complexes critical for stem cell function including the PRC2, BAF and NuRD complexes. stem.MAP also discriminates mutually exclusive interactions specific to embryonic stem cells (e.g. Eif4ebp2 - Eif4e). Finally we observe 1,388 high confidence stem.MAP interactions found only in mESC CF-MS data, many with support from AlphaFold modeling and a gastruloid coregulation network. We make stem.MAP protein complexes available through our web resource: stem.proteincomplexes.org. We anticipate stem.MAP will be beneficial to the study of early development and we expect our proteomics-based machine learning workflow to be applicable to the creation of any context-specific protein interaction network.

## Introduction

Proteins assemble to form stable macromolecular protein complexes, which carry out cellular function and determine cellular fate. For the past two decades, major efforts have leveraged high-throughput proteomics to identify protein complexes in various organisms including human (Hein *et al*, 2015; Huttlin *et al*, 2021, 2017, 2015; Wan *et al*, 2015), yeast (Gavin *et al*, 2002; Krogan *et al*, 2006), and plants (McWhite *et al*, 2020). Manual literature curation has also increased the coverage of protein complexes across species (Giurgiu *et al*, 2019; Balu *et al*, 2025). More recent efforts have focused on integrative approaches using machine learning to identify protein complexes across thousands of proteomic experiments (Drew *et al*, 2017, 2021; Fischer *et al*, 2025). While these efforts have successfully increased coverage and accuracy of identifying protein complexes, only a subset is likely assembled in a given cell type or tissue at a given time.

Early studies identified context-specific interactions by overlaying gene expression data onto protein interaction networks. A study in yeast, for example, showed protein interactions were dynamic across the cell cycle, highlighting the condition-specific nature of complexes (de Lichtenberg *et al*, 2005). In human tissue, gene expression was similarly used to infer that universally expressed proteins tended to make many interactions with tissue-specific proteins, further demonstrating the importance of tissue-specific interactomes across distinct tissues (Bossi & Lehner, 2009). Coupling gene expression with protein interactions is valuable but unfortunately masks complexes that assemble independent of gene expression. More recent work has applied high-throughput proteomics assays including affinity purification mass spectrometry (AP-MS) to specific cell types and tissues (Huttlin *et al*, 2021). These efforts provide direct evidence of cell-type-specific interactions instead of inferring them through gene expression based approaches (Magger *et al*, 2012; Ziv *et al*, 2022). For example, analyses of the BioPlex3 HEK293T and HCT116 interaction networks found that only 50% interactions were shared between the cell lines, indicating a substantial rewiring of protein networks between them (Huttlin *et al*, 2021).

Complementary methods such as co-fractionation mass spectrometry (CF-MS) have been particularly successful in exploring stable protein complexes across the proteome (Wan *et al*, 2015; Cox *et al*, 2026; Goel *et al*, 2024) as well as in specific tissues, including red blood cells and neurons (Sae-Lee *et al*, 2022; Skinnider *et al*, 2021; Dang *et al*, 2026). CF-MS involves biochemical fractionation of non-denatured lysate followed by mass spectrometry and identifies macromolecular complexes by observing co-eluting proteins across many orthogonal separation techniques (e.g. size exclusion chromatography (SEC), ion exchange chromatography (IEX)). CF-MS is label-free, requiring no genetic tags or antibodies, theoretically allowing application to any cell type or tissue. CF-MS easily scales to the entire proteome but requires a substantial number of mass spectrometry experiments to confidently identify protein complexes.

Here, we propose a prior-based strategy to predict protein complexes in mouse embryonic stem cells (mESCs) using <250 mass spectrometry experiments, an order of magnitude less than other cell-type-specific protein complex maps (Huttlin *et al*, 2021; Sae-Lee *et al*, 2022). mESCs are both self-renewing and pluripotent, capable of differentiating into all other cell types (Martello & Smith, 2014). The molecular basis for these functions, especially at the interactome level, remain underexplored. Identifying protein complexes specific to mESCs offer direct molecular insights into their roles in self-renewal and pluripotency. Here, we construct a machine learning pipeline which integrates CF-MS data from mESCs with prior information from global interaction networks to build a cell-type-specific protein complex map (stem.MAP). Our map identifies 3,986 protein complexes in embryonic stem cells involving 8,616 unique proteins. We achieve high precision and recall by incorporating priors from a global protein complex map (i.e. hu.MAP3.0) and several pre-training steps to ensure identification of cell-type-specific complexes. We highlight several stem cell-specific complexes associated with human disease and identify novel stem cell-specific protein interactions. Specifically, we identify known stem cell-specific components of chromatin remodelers including Jarid2, Mtf2, Aebp2 and Epop in the PRC2 complex, SmarcA4, Dpf2, Arid1a, SmarcC1 and SmarcD1/D2 in the BAF complex, and Wdr5 in the NuRD complex. We also identify 1,388 protein interactions only found in mESC CF-MS data including interactions involved in protein translation, cell cycle regulation, and lipid metabolism necessary for ES expansion. Of these, 28 are supported by AlphaFold3 modeling and 63 are supported by the gastruloid co-regulation network (Garge *et al*, 2026). Finally, we made a publicly accessible interface to all stem.MAP protein complexes (stem.proteincomplexes.org).

## Results

### Cell-type-specific protein complex map machine learning workflow

Embryonic stem cells are a critical cell type which hold the potential to differentiate into all other cell types (Martello & Smith, 2014). Core transcription factors (e.g. Oct4, Sox2, Klf4, and c-Myc) are important for these functions (Takahashi & Yamanaka, 2006) but other factors such as chromatin remodelers also play a role. The BAF complex, for example, is a chromatin remodeler that functions during embryonic development and consists of different subunits depending on the context (Zhang *et al*, 2019). Global protein complex maps such as hu.MAP (Fischer *et al*, 2025; Drew *et al*, 2017, 2021) do not characterize complexes in a cell-type-specific manner (Figure 1A) and therefore lack context on BAF complex variants. Though studies have explored protein interaction networks in stem cells, they remain restricted to core transcription factors (Pardo *et al*, 2010; van den Berg *et al*, 2010; Wang *et al*, 2006). We still lack a systematic understanding of the protein complexes and their variants present in stem cells.

**Figure 1.**
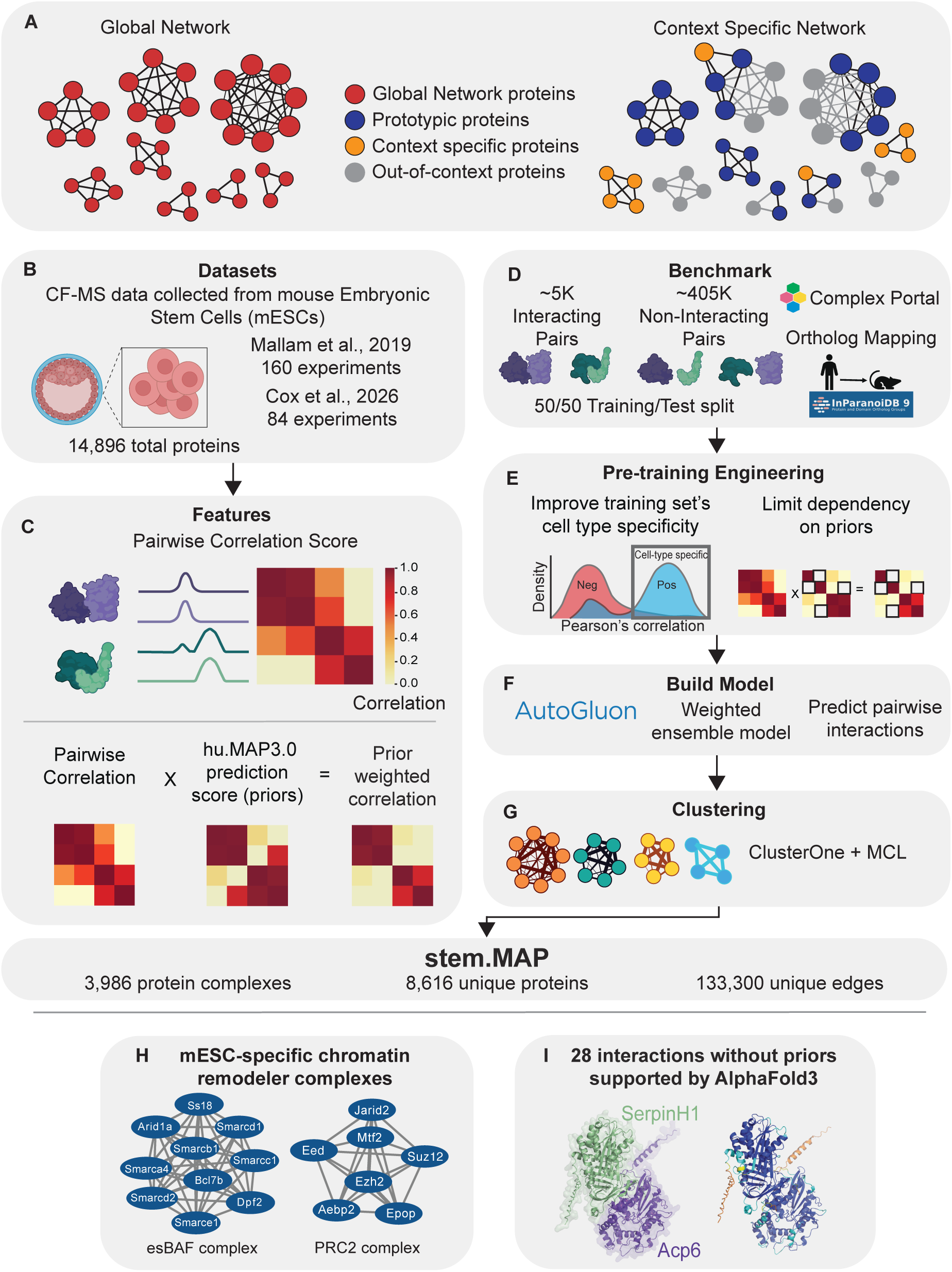
stem.MAP Workflow. **A**. Global network vs context specific network. A global network identifies protein interactions but does not consider interactions in different cellular contexts. A context specific network identifies interactions in different cellular contexts. Prototypic proteins are subunits found in multiple contexts. Context specific proteins are subunits found primarily in the context of interest. Out-of-context proteins are subunits not found in the context of interest. **B.** Co-Fractionation Mass Spectrometry (CF-MS) data from mouse embryonic stem cells were collected from Mallam et al., 2019 and Cox et al., 2026. **C.** Pearson correlation coefficient was calculated for all pairs of proteins. Pairwise correlation scores were multiplied by huMAP3.0 confidence scores to generate prior-weighted features. **D.** Human protein complexes from ComplexPortal were ortholog-mapped to mouse protein IDs. Interacting pairs (positives) are present in the same complex, non-interacting pairs (negatives) are proteins in separate complexes. **E.** Pre-training engineering improved cell-type-specificity of the training benchmark and reduced reliance on prior-based features. **F.** AutoGluon Tabular was used as a machine learning classification framework to build a weighted ensemble model and predict pairwise interactions between proteins. **G.** A two-stage clustering was used to identify protein complexes within the generated protein interaction network. **H.** stem.MAP captures mESC-specific variants of chromatin remodeler complexes. **I.** stem.MAP predicts novel cell-type-specific protein interactions which are supported by AlphaFold3 modeling. *Figure partially created with BioRender.com*

To construct a cell-type-specific protein complex map for embryonic stem cells, we developed a proteomics-based machine learning workflow (Figure 1B-G) to process CF-MS data from mESC. We collected three CF-MS experiments on mESCs, including two size exclusion chromatography (SEC) experiments (Mallam *et al*, 2019) and one ion exchange (IEX) chromatography experiment (Cox *et al*, 2026) (Figure 1B). All three experiments identify ∼15k proteins (>= 2 PSMs, see Methods), providing substantial coverage of the mESC proteome.

To identify co-eluting proteins that suggest co-complex interactions, we calculated a similarity value for all protein pairs in each experiment based on Pearson correlation (Figure 1C, see Methods) (McWhite *et al*, 2021). Protein complex maps built using CF-MS experiments typically rely on several thousands of fractions, constituting tens of chromatography experiments. Given that we only had access to three high-quality mESC CF-MS datasets, we constructed a framework that incorporates previously identified protein interactions from hu.MAP3.0 to be used as priors. We created three additional features based on the correlation scores calculated from these mESC CF-MS datasets multiplied by hu.MAP3.0 pairwise confidence scores (Figure 1C). Our rationale for using prior-weighted features was that protein pairs with spuriously high correlation values calculated from the mESC data would be downweighted if limited evidence was previously seen (i.e. low confidence score) in hu.MAP3.0. Alternatively, protein pairs with lower correlation values from mESC data could potentially be rescued by high confidence scores in hu.MAP3.0. Both of these scenarios rely on protein pairs being scored by hu.MAP3.0, which would preclude any mESC-specific protein interactions. In our workflow, we therefore include unweighted correlation features in addition to our prior-weighted features to allow for the identification of protein interactions found only in stem cells.

Creating a cell-type-specific protein complex map using a machine learning workflow requires training and evaluating on a benchmark of protein pairs known to interact (positives) as well as pairs that do not (negatives). Due to the scarcity of known mESC-specific interactions in curated databases, we rely on a cell-type-agnostic benchmark of protein complexes rather than a cell-type-specific benchmark. Importantly, this allows our workflow to be applied to nearly any cell type. To avoid data leakage within our machine learning benchmark due to our use of hu.MAP3.0 priors, we used the same test and training benchmark split of ComplexPortal complexes(Balu *et al*, 2025) used by hu.MAP3.0. This resulted in a total of ∼5,000 positive interacting protein pairs, and ∼405,000 non-interacting protein pairs (Figure 1D, see Methods).

Using a cell-type-agnostic benchmark introduces its own challenges that need to be addressed prior to training a machine learning model. First, known positive protein interactions in our benchmark may not be present in the available CF-MS data and could potentially skew the training, making it difficult to discriminate between true and false interactors. We therefore calculate the correlation score for all positive training protein pairs in the mESC CF-MS data and remove the positive pairs from the training set with low correlation values (Figure 1E, see Methods). Pairs with low correlation values in cell-type-specific CF-MS data are likely interactions not present in the cell type and therefore are not trained on. Importantly, the test set is not altered and remains a true evaluation of the performance of the model.

Another challenge when training a cell-type-specific model with a cell-type-agnostic benchmark arises from the high prevalence of benchmark protein pairs having been studied previously. Specifically, all protein pairs in our benchmark set have prior information while only ∼20% of protein pairs in our CF-MS data have prior information (i.e. ∼18 million pairs with priors out of 96 million pairs with CF-MS data). This skews the importance of prior-weighted features during training and eliminates the ability to identify real cell-type-specific protein interactions. We addressed this by removing prior information for a randomly selected 10% of training pairs to allow our model to identify protein pairs without prior information (Figure 1E, see Methods).

Next, we use an automated model selection algorithm, Autogluon (Erickson *et al*, 2020), to train a machine learning classifier to discriminate between interacting (i.e. co-complexing) vs. non-interacting protein pairs (Figure 1F). Autogluon first builds 13 different models and then constructs an ensemble model based on the highest performing original models (Supplemental Table 1). The final weighted ensemble model consisted of LightGBM (0.66) and NeuralNetTorch (0.33). This final model was then applied to all protein pairs in the mESC CF-MS dataset to generate a confidence score of their likelihood of interacting. We cluster the resulting all-by-all protein interaction network to identify highly connected regions representing protein complexes (Figure 1G, see Methods). The final network, stem.MAP, identifies a total of 3,986 protein complexes (Supplemental Table 2) consisting of 8,616 unique proteins, and 133,300 unique protein interactions (Supplemental Table 3). In addition to identifying globally expressed protein complexes (e.g. proteasome, ribosome), stem.MAP identifies stem cell-specific protein complexes including several disease-relevant chromatin remodeler complexes (Deevy & Bracken, 2019; Valencia *et al*, 2023) (Figure 1H). Additionally, stem.MAP identifies 1,388 stem cell-specific protein interactions with no prior information, 28 of which are supported by AlphaFold modeling (Figure 1I). These represent novel interactions that potentially are found only in the stem cell state.

### Incorporation of ‘priors’ and pre-training engineering improves the prediction accuracy of stem.MAP

As mentioned above, we needed to address several challenges when using a cell-type-agnostic benchmark to train and evaluate a cell-type-specific model. We removed positive training examples which are unlikely to be present in mESC (Figure 2A). To do this, we calculated Pearson correlation coefficients for all positive and negative training elution profiles from the three mESC CF-MS datasets and averaged the correlation coefficients. We observed a considerable density of positive pairs with low average correlations that overlapped with the expected low average correlations of negative pairs (Figure 2B). We calculated the percentage of positive pairs below a range of thresholds and chose a conservative threshold (< 0.1 average correlation, Supplemental Figure 1). In total, we removed 1,035 training pairs below the threshold, leaving 58% (i.e. 1,452) positive pairs with data for training. To ask how the removal of these training pairs affected the performance of our model, we trained one model on the full training set and another model on the training set with low-correlation positives removed. The model with removed training examples shows higher precision for high-confidence predictions representing a better overall model (Figure 2C).

**Figure 2.**
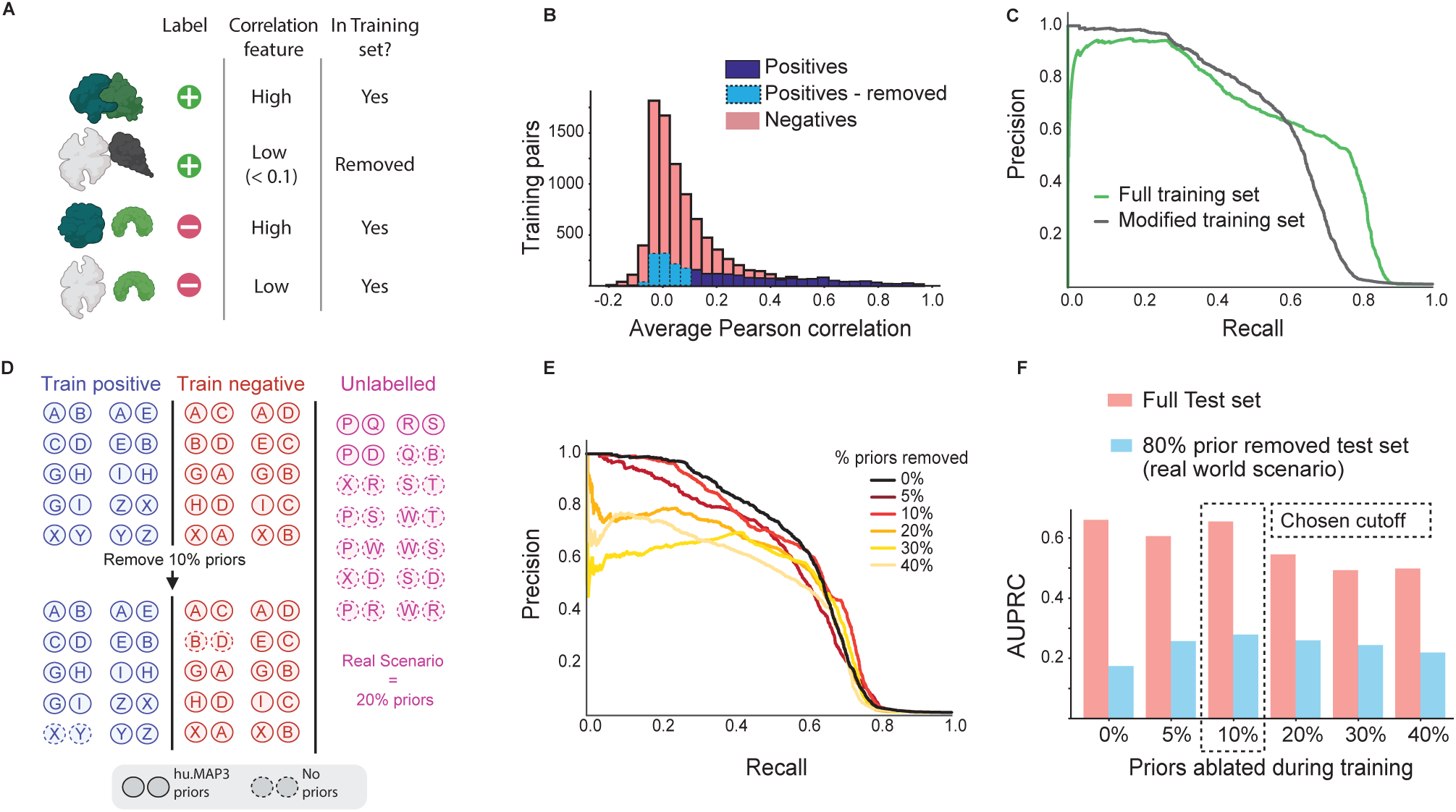
Pre-training engineering increases the context specificity of the training benchmark. **A**. Due to the training set having positive examples that do not exist in mESCs, positive training examples with low pairwise Pearson correlation coefficient were removed. The test set was unaltered. **B.** Histogram of average Pearson correlation scores across three CF-MS datasets of positive (blue) and negative (red) training labels. A cutoff correlation value of 0.1 was used to remove 1,035 low correlation positive training examples (light blue, dashed outline). **C**. Precision-recall curves evaluating classification models on the leave-out test set before (green) and after (black) removal of training positives (modified training set). **D.** Schematic of our strategy to train and evaluate our model to recognize protein pairs without prior data. To train, 10% of prior data for training examples was ablated from both the positive and negative pairs. To evaluate, 80% of prior data of the test set was ablated to mimic the real world scenario where most protein pairs lack prior data in the hu.MAP3.0 protein interaction network **E.** Precision-recall curves evaluating classification models with different amounts of prior data ablated from the training set. The evaluation is done on the full test set with no priors ablated. The model with 10% training prior data ablation (light red) shows minimal loss in performance compared to the full prior data model (0% removal, black). **F.** Bar plot shows the area under the precision recall curve (AUPRC) for models with different amounts of prior data ablated during training, evaluated with the full test set (red), and a test set with 80% prior data ablated (blue) mimicking the real world scenario. The model with 10% prior data ablated performs comparably to the top (0% ablated) model in the full test set and outperforms all models in the real world scenario. *Figure partially created with BioRender.com*

We observed that all training and testing examples in our benchmark have prior information (i.e. hu.MAP3.0 confidence scores). This poorly represents a real-world scenario where only ∼20% of protein pairs with CF-MS data have prior information in hu.MAP3.0. Training and testing on a benchmark with complete prior information prevents the model from identifying protein interactions not previously seen in hu.MAP3.0. To address this issue, we removed prior information for a randomly selected set of benchmark proteins (Figure 2D). We trained several models on sets with different percentages of priors removed and compared their performances on the unaltered leave-out test set. We observe a substantial drop in performance (-0.1 AUPRC) when removing greater than 10% of priors. Further, removing 10% had minimal impact on performance compared to the full set, AUPRC = 0.594 vs 0.598 respectively (Figure 2E). When comparing each of these models against a real-world scenario where 80% of the pairs in the test set have no hu.MAP3.0 priors, we found that the model trained with 10% of the priors removed performs best (Figure 2F). We therefore used the 10% prior cutoff for our training going forward.

### stem.MAP is highly accurate and recovers embryonic stem cell-specific biology

We next evaluate the contribution of each individual feature type to the overall performance of the final model. We found that prior-weighted features perform substantially better than individual correlation features (Supplemental Figure 2A, Figure 3A, see Methods). Although priors add more recall, we observed that our model’s highest confidence predictions were driven by the Pearson correlation features (Figure 3A). Moreover, we see the prior-weighted correlation-only model performs well overall but lacks precision in the high-confidence region (Recall < ∼0.3). This early performance drop is recovered by the full model when unweighted correlation values are included (Figure 3A), indicating both feature types add value to the total performance of the final model. We evaluated the calibration of our model by plotting the relationship between our confidence thresholds and the precision on the leave-out test set (Supplemental Figure 2B). We see our model is slightly overconfident for the high-confidence thresholds and underconfident for the lower thresholds.

**Figure 3.**
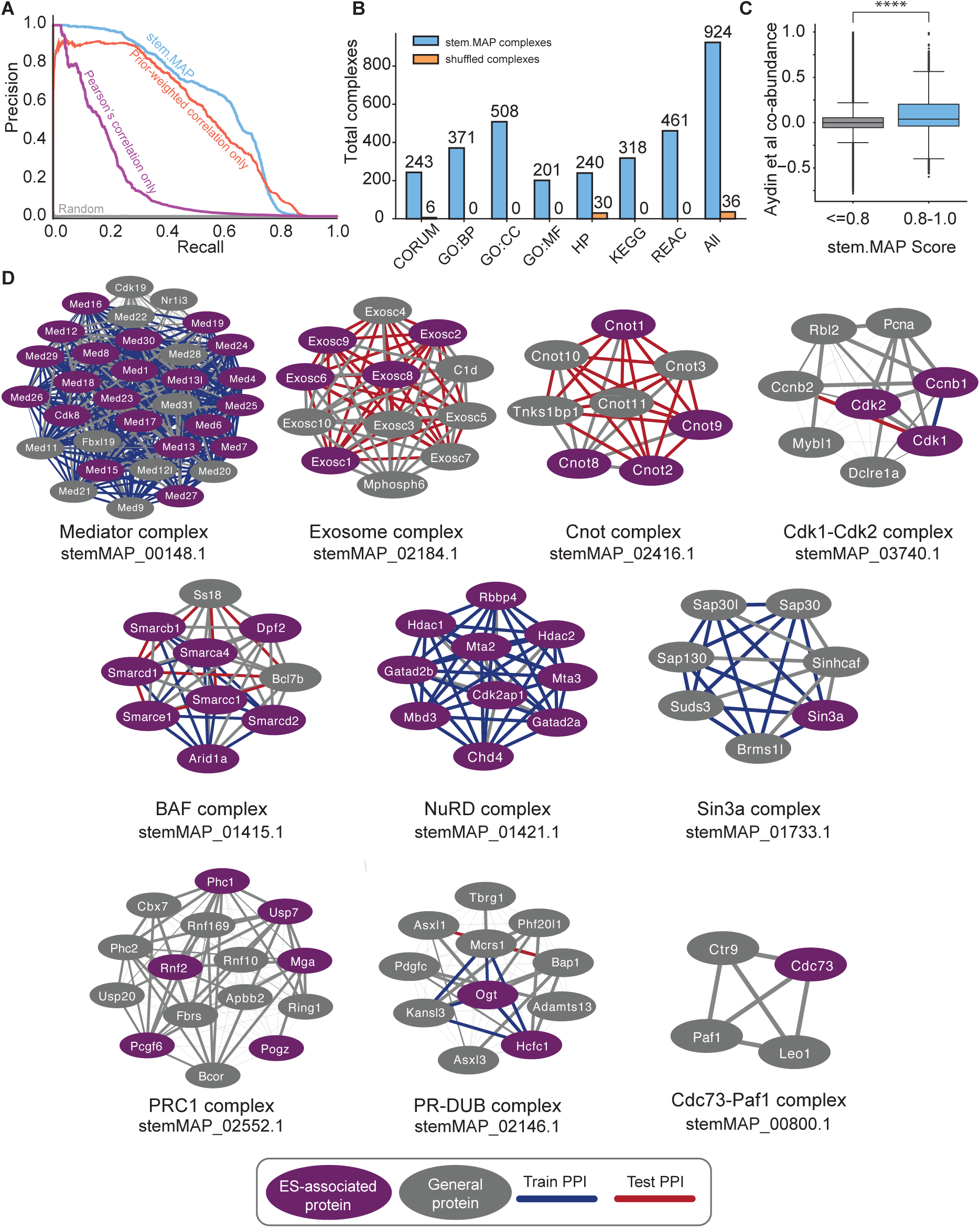
Global evaluation of stem.MAP. **A.** Precision recall curve of stem.MAP weighted ensemble model predictions using all features (blue) compared with Pearson correlation score features only (prior-weighted features ablated, purple) and prior-weighted correlation features only (correlation only features ablated, red). **B.** Bar graph shows annotation enrichment of stem.MAP predicted complexes (blue) for seven annotation types. Annotation enrichment for complexes is determined by g:Profiler with a p-value cutoff of 0.01. Subunits of stem.MAP protein complexes were shuffled to determine annotation enrichment by chance (blue). **C.** Protein pair co-abundance values calculated from genetically diverse mouse embryonic stem cells show statistically different distributions between low scoring (< 0.8) and high scoring (>= 0.8) stem.MAP protein interactions (two tailed KS test, statistic=0.2, pvalue=0.0, **** = p-value <= 0.0001). **D.** stem.MAP identified protein complexes relevant to embryonic stem cells such as the mediator, exosome and Cnot1 complex, cell cycle regulators like the Cdk1-Cdk2 complex, and chromatin modifier complexes like BAF complex, NuRD complex, Sin3a complex, PRC1 complex, PR-Dub complex and Paf1 complex. stem.MAP IDs of the complex are listed below each complex. ES-associated proteins (purple) are from (Ng & Lufkin, 2011; Van Hoof *et al*, 2006; Atlasi *et al*, 2020). Train (blue) and Test (red) edges are highlighted. Edge weights are proportional to stem.MAP confidence score.

We next apply a two-stage clustering pipeline to the full protein interaction network generated by our machine learning classifier. We optimized our clustering pipeline by sweeping clustering parameters and comparing each resulting clustering to a set of literature curated training complexes. As has been seen previously (Drew *et al*, 2017, 2021; Fischer *et al*, 2025), optimal clusterings are a tradeoff between complex-level precision and recall resulting in a Pareto front (Supplemental Figure 2C). We therefore selected two parameter combinations which represent a high-precision - low-recall clustering and a high-recall - low-precision clustering. Complexes in the high-precision clustering tend to be smaller on average, with 84% of clusters containing < 5 subunits, whereas only 22% of complexes in the high-recall clustering consisted of < 5 subunits (Supplemental Figure 2D). Evaluating the high-precision, high-recall, and a unified set of both high precision and high recall against the literature-curated leave-out test complexes (Supplemental Figure 2E), we observe consistent performance between the test and training complexes suggesting our model and clustering is generalized in identifying protein complexes.

We next asked if the stem.MAP complexes produced by our machine learning pipeline capture known biology in the literature. We anticipated that proteins in the same complex should share biological functions, including similar subcellular localizations. We used g:Profiler (Kolberg *et al*, 2023) to identify enriched annotations (p value <= 0.01) for each stem.MAP complex (Figure 3B). We identified 924 stem.MAP complexes with at least one enriched annotation, and hundreds of complexes enriched for GO (Ashburner *et al*, 2000) Biological Process, Cellular Component, Molecular Function, Human Phenotype Ontology (Köhler *et al*, 2014) terms, KEGG (Kanehisa *et al*, 2014) terms, Reactome (Milacic *et al*, 2024) terms, and CORUM complexes (Giurgiu *et al*, 2019) (see Supplemental Table 4). To control for complexes being associated with annotations by chance, we ran the same analysis on randomly shuffled complexes and observed only 36 shuffled complexes with enriched annotations, demonstrating the biological relevance of stem.MAP complexes.

To understand if stem.MAP is highly correlated in abundance across multiple experiments, we compared stem.MAP interactions to proteomics co-abundance values calculated from 190 genetically outbred mESC cell lines (Aydin *et al*, 2023). Specifically, we calculated pairwise Pearson correlation coefficients (co-abundance scores) from normalized protein abundance values across the 190 mESC cell lines. Of the ∼17 million pairs of proteins with both co-abundance and stem.MAP scores, 98,257 pairs with stem.MAP score >= 0.8 had significantly higher co-abundance compared to lower stem.MAP scoring pairs (Figure 3C, KS test, p = 0.0). For example, we see high-confidence stem.MAP predictions between subunits of the mitochondrial ribosome having high co-abundance values (Mrpl4 and Mrpl40, stem.MAP score = 0.98, co-abundance value = 0.99, Supplemental Table 5). On the other hand, we observe Jarid2 and Suz12, two proteins associated with the Polycomb repressive complex 2 (PRC2) important for pluripotency (Lackner *et al*, 2021), having high-confidence stem.MAP predictions and low co-abundance values (stem.MAP score = 0.99, co-abundance value =-0.05). Jarid2 is an accessory subunit to the PRC2 complex and is only present in a subset of PRC2 complexes (Shen *et al*, 2009), explaining the low co-abundance value.

A core set of transcription factors have been identified as regulators of embryonic stem cell self-renewal and pluripotency including Klf4, Nanog, Oct4/Pou5f1, Myc, and Sox2 (Okita *et al*, 2007; Takahashi & Yamanaka, 2006). Previous studies have demonstrated these transcription factors physically interact with each other as well as the other transcriptional machinery (Wei *et al*, 2009; Pardo *et al*, 2010). We observe Klf4, Nanog, Oct4/Pou5f1, and Sox2 coelute in the CF-MS SEC experiment consistent with their previously described physical interactions (Supplemental Figure 3A). stem.MAP identifies these factors in several protein complexes with previously unassociated proteins. For example, Oct4/Pou5f1 is identified in a large complex with 18 subunits (Supplemental Figure 3B, stemMAP_00115.1). Among the members, L1td1 is known to interact with Pou5f1 (Emani *et al*, 2015) and several of the subunits, L1td1 and Utf1 (Okuda *et al*, 1998), are known ES-associated proteins. Interestingly, none of the interactions involving Pou5f1 had priors from hu.MAP3.0 suggesting this is an ES-specific complex. Additionally, stem.MAP identifies a complex related to genome integrity consisting of subunits Klf4, Tp53bp1, Slf2, and Psrc1 (Supplemental Figure 3C, stemMAP_01109.1). None of the interactions involving Klf4 have priors from hu.MAP3.0 suggesting this is also an ES-specific complex.

We next compare the proteome coverage in stem.MAP to other large-scale proteomics datasets from mouse embryonic stem cells (Van Hoof *et al*, 2006; Atlasi *et al*, 2020). Van Hoof et al. collected proteomics data on embryonic stem cells and differentiated cells, identifying 743 and 424 unique proteins within each respectively. Additionally, Atlasi et al. identified ∼3,900 unique abundant proteins from naïve embryonic stem cells. stem.MAP has considerable overlap with Atlasi-specific proteins (i.e. 2,495 overlapping proteins) as well as the intersection of Atlasi and Van Hoof mESC-unique proteins (i.e. 402 overlapping proteins) (Supplemental Figure 3D). We observed many stem.MAP protein complexes consist of multiple ES-associated proteins found in Atlasi and Van Hoof mESC datasets, including the mediator (e.g. Med1, Med12), BAF (e.g. Arid1a, Smarca4), NuRD (e.g. Mbd3, Hdac1), and PRC1 (e.g. Pcgf6, Mga) complexes (Figure 3D). Only 44 proteins were not recovered by stem.MAP that are in both other datasets, suggesting stem.MAP has high coverage of the mESC proteome. We also see our CF-MS datasets show high abundance for mESC-specific proteins and lower abundance for differentiation-specific proteins identified by Van Hoof demonstrating the cell type specific nature of stem.MAP (Supplemental Figure 3E).

### stem.MAP identifies known embryonic stem cell-specific chromatin remodeler complexes

Embryonic stem cells rely on transcription factors interacting with different chromatin remodeler complexes to maintain pluripotency (Golob *et al*, 2008). Using stem.MAP, we identified stem cell-specific variants of known chromatin remodeler complexes. Here, we specifically focus on chromatin remodeler complexes where certain variants of the complex are known to interact in ES cells but not differentiated cells. The PRC2 complex, for example, is a known chromatin remodeler which methylates chromatin into a repressive state and is important for maintaining pluripotency (Margueron & Reinberg, 2011). Several PRC2 subunits are specific to ES cells including Jarid2, Aebp2, Ezh2, while other subunits have been identified in many cell types, which are termed here as prototypic subunits (Figure 4A). Our pipeline identifies a PRC2 complex which includes both Prototypic and ES subunits (Figure 4B, stemMAP_01662.1). Elution profiles for PRC2 related proteins show strong co-elution of many of the ES subunits with the Prototypic subunits and low co-elution of non-ES subunits (e.g. Ehmt2 and Phf1) (Figure 4C).

**Figure 4.**
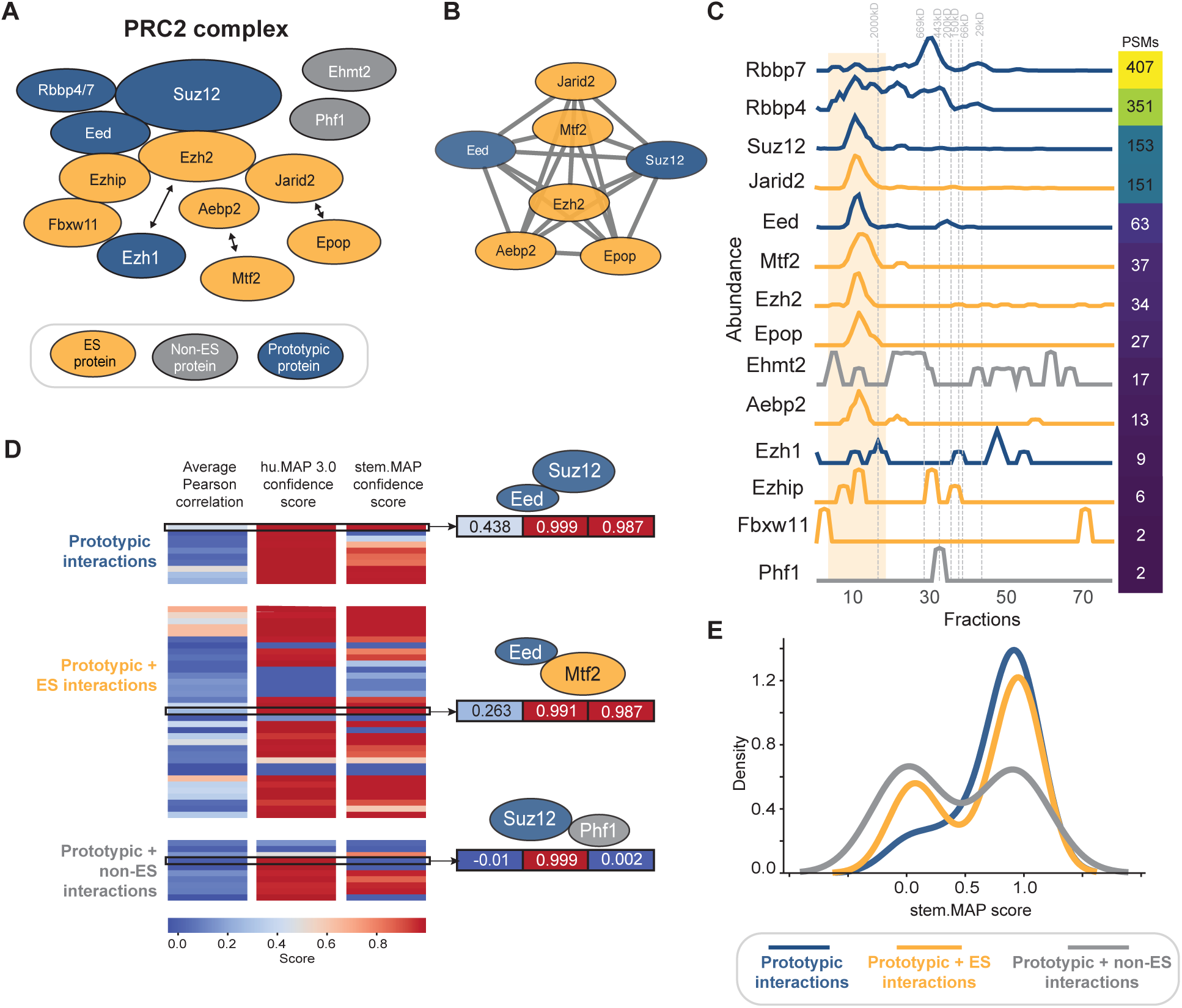
stem.MAP predicts stem cell-specific interactions. **A.** Literature curated PRC2 complex. Blue nodes represent proteins in the complex that are shared between different cell types (Prototypic proteins). Yellow nodes represent proteins that are specific to embryonic stem cells or necessary for ES-self renewal and pluripotency (ES proteins). Gray nodes represent interactors of the complex not present in embryonic stem cells (Non-ES proteins). Bidirectional arrows indicate mutually exclusive subunits. **B.** PRC2 complex identified by stem.MAP (stemMAP_01662.1). **C**. Elution profiles of PRC2 associated subunits from mouse embryonic stem cell SEC CF-MS experiment. Subunit colors match that described in **A.** Profiles are normalized to the highest abundant fraction for each subunit. Total abundance values (peptide spectral matches = PSM) are listed for each subunit. The highlighted region (yellow) represents fractions where the PRC2 complex is present. **D**. Colored heatmap of average Pearson correlation coefficient, hu.MAP3.0 confidence score (i.e. prior), and stem.MAP confidence score for pairs consisting of proteins in literature curated PRC2 complex. Pairs are classified into Prototypic interactions (blue), Prototypic + ES interactions (yellow), and Prototypic + non-ES interactions (gray). Specific examples of each category are highlighted to the right of the heatmap. **E**. Distributions of stem.MAP confidence scores for each classification in **D**. Prototypic and Prototypic + ES interactions are right shifted (i.e. more confident) while Prototypic + non-ES interactions are left shifted (i.e. less confident).

To determine if our model can discriminate between ES and non-ES subunits, we created three classes of interactions in the PRC2 complex: 1) interactions among the Prototypic subunits (Prototypic interactions), 2) interactions between Prototypic subunits and ES subunits (Prototypic + ES interactions), and 3) interactions between Prototypic subunits and non-ES subunits (Prototypic + non-ES interactions). We then evaluated interactions in each class based on the correlation of mESC CF-MS data, the prior values (i.e. hu.MAP3.0 confidence scores), and the final stem.MAP confidence score. We represent scores of all pairs in each class as a heatmap (Figure 4D). A representative Prototypic interaction example is Eed and Suz12 with partial correlation in mESC CF-MS data (0.438) and a high prior (0.999). The stem.MAP confidence score results in a strong prediction (0.987) as expected for the Prototypic subunits. Further, we observe the Prototypic + ES interaction, Eed and Mtf2, having partial correlation (0.263) in the CF-MS data and a high prior (0.991) resulting in a strong prediction interaction by stem.MAP (0.987). This suggests stem.MAP can identify ES interactions even among subunits with partial correlation. Finally, we observe no correlation (-0.01) in the CF-MS data for the Prototypic + non-ES interaction (Suz12 and Phf1) even though the pair has a strong prior (0.999). The resulting stem.MAP confidence score for Suz12 - Phf1 has very low confidence (0.002). Comparing the overall distributions of these classes, we see Prototypic interactions and Prototypic + ES interactions have high stem.MAP scores while Prototypic + non-ES interactions have lower stem.MAP scores (Figure 4E).

Other ES protein complexes exhibit a similar trend. Both the BAF and NuRD complexes are critical for ES function of pluripotency and self-renewal (Reynolds *et al*, 2012; Ho *et al*, 2009) with each having their own ES components. We see ES subunits of the BAF complex including Arid1a, Smarcc1, and Smarcd1 (Figure 3D, Supplemental Figure 4A) and importantly do not see non-ES subunits such as Arid1b, Smarcc2, or Smarcd3. Similar to the PRC2 complex, stem.MAP discriminates between BAF Prototypic + ES interactions (high stem.MAP scores) and Prototypic + non-ES interactions (low stem.MAP scores) (Supplemental Figure 4B-C). This is consistent with the elution profiles from the CF-MS data for the BAF subunits (Supplemental Figure 5). Regarding the NuRD complex, stem.MAP identifies a critical protein required for pluripotency exit, Mbd3, as well as the ES subunit, Wdr5 as components of the NuRD complex and, importantly, do not see the non-ES subunits Bcl11b or Chd5 (Figure 3D, Supplemental Figure 4D). The majority of Prototypic interactions and the Prototypic + ES interactions of the NuRD complex have high stem.MAP confidence scores while the Prototypic + non-ES interactions have mostly low stem.MAP confidence scores (Supplemental Figure 4E&F). This is supported by the elution profiles from CF-MS data for NuRD subunits (Supplemental Figure 6). Overall, stem.MAP appropriately discriminates between ES and non-ES subunits for these complexes reinforcing the idea that it identifies protein complexes critical for ES function.

### stem.MAP prefers one mutually exclusive interactor over another

The functions of protein complexes can change depending on individual subunits swapping into or out of a complex based on different cellular conditions or cell types. Examples include the catalytic activity of the proteasome (PSMB5 <-> PSMB8), the transcriptional output of RNA Pol III (POLR3G <-> POLR3GL), and the chromatin remodeling of the BAF complex (ARID1A <-> ARID2). We recently used structural modeling to identify > 5,800 mutually exclusive protein pairs in hu.MAP3.0 (Fischer *et al*, 2025). We hypothesized that many mutually exclusive pairs would show preference for one member over the other in embryonic stem cells (Figure 5A). We therefore asked if stem.MAP shows a confidence score discrepancy between the two proteins and their common subunit. For all mutually exclusive proteins identified in hu.MAP3.0 whose orthologs were also scored by stem.MAP, we calculate a Confidence Score Delta (see Methods), which is the magnitude of the difference in confidence scores between two mutually exclusive protein pairs sharing a common subunit (Figure 5A). We compared the calculated Confidence Score Delta of mutually exclusive proteins as predicted by hu.MAP3.0 and stem.MAP (Figure 5B, Supplemental Table 6). As expected by the cell-type-agnostic nature of the network, hu.MAP3.0 reported similar confidence scores between mutually exclusive subunits and their common subunit. Alternatively, stem.MAP predictions report large differences in confidence scores between mutually exclusive proteins and their common subunits, indicating one protein may be preferred over the other in embryonic stem cells.

**Figure 5.**
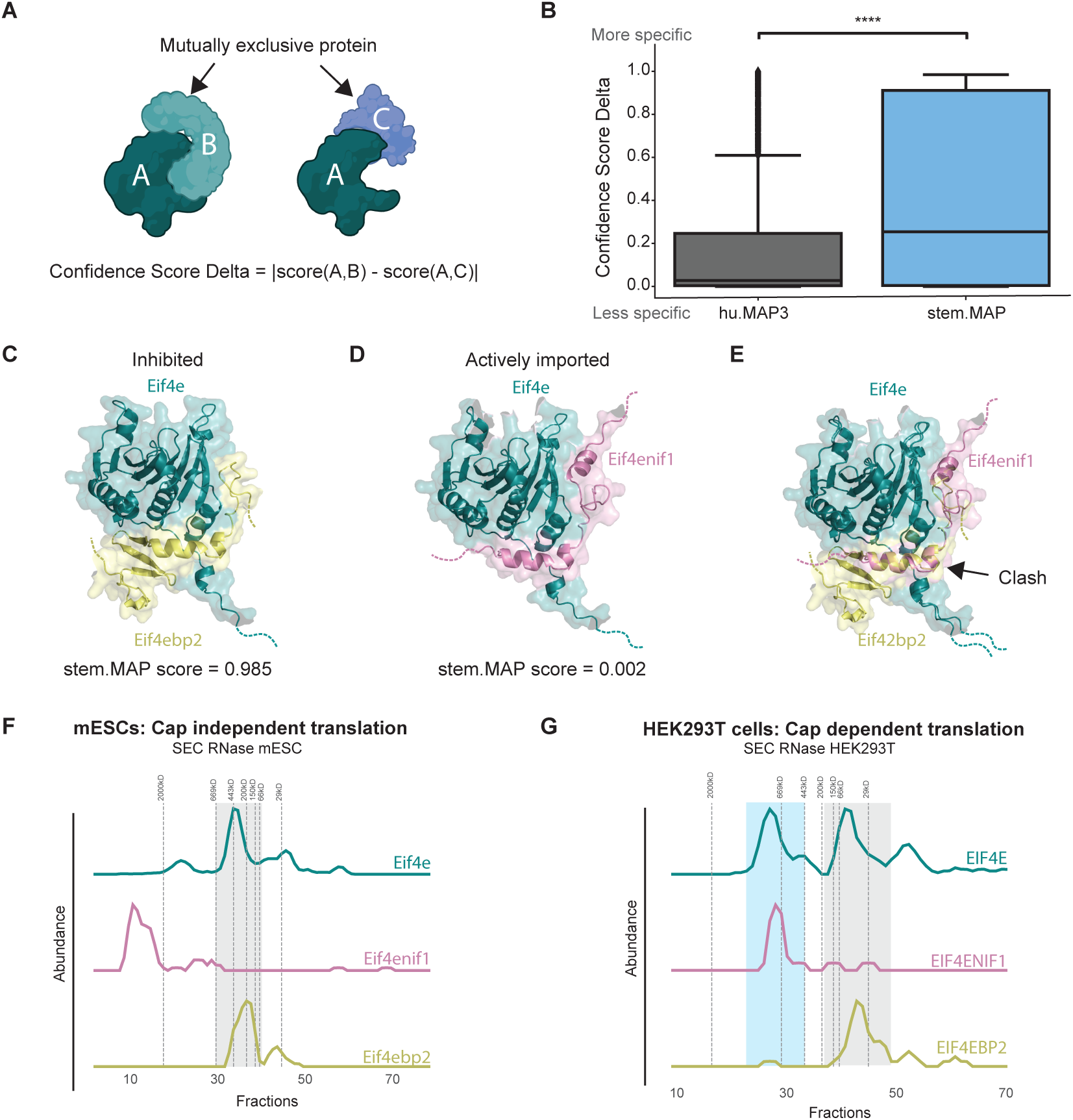
stem.MAP differentiates between preferred partners in a mutually exclusive interaction. **A.** Illustration of mutually exclusive interactions; proteins B and C share the same structural interface with a common subunit A. Confidence Score Delta is the difference in prediction scores between each mutually exclusive interaction with the common subunit. **B.** Distributions of Confidence Score Deltas of mutually exclusive proteins for hu.MAP3.0 (gray) and stem.MAP (blue). The context specific stem.MAP network confidently identifies one specific mutually exclusive subunit over another while the context agnostic hu.MAP3.0 network scores mutually exclusive subunits similarly (**** p-value = 1.3e-70, Mann–Whitney U test). **C.** Structural model of interaction between Eif4e and Eif4ebp2. stem.MAP confidence score = 0.985, hu.MAP3.0 confidence score = 0.998. **D.** Structural model of Eif4e and Eif4enif1. stem.MAP confidence score = 0.002, hu.MAP3.0 confidence score = 0.999. **E.** Overlay of structural models from **C** and **D**. Mutually exclusive subunits Eif4ebp2 and Eif4enif1 share an interface when interacting with Eif4e which would result in a clash. **F.** Elution profiles of Eif4e, Eif4enif1, and Eif4ebp2 from a size exclusion chromatography CF-MS experiment on mouse embryonic stem cells. Eif4e and Eifebp2 coelute (gray highlight) while Eif4enif1 elutes separately. This represents an inhibited Eif4e complex and denotes an absence of the actively imported complex in the nucleus. Elution profiles of EIF4E, EIF4ENIF1, and EIF4EBP2 from a size exclusion chromatography CF-MS experiment on HEK293T cells. EIF4E separately coelutes with both EIF4ENIF1 (blue highlight) and EIF4EBP2 (gray highlight) suggesting both interactions are formed but in separate contexts.

For example, Eif4ebp2 and Eif4enif1 both interact with Eif4e. Eif4e is a mRNA 5’ cap-binding protein critical for translation initiation (Yang *et al*, 2020) and is involved in mRNA export from the nucleus (Culjkovic *et al*, 2006). Eif4enif1 facilitates the localization of Eif4e to the nucleus (Dostie *et al*, 2000). Eif4ebp2, on the other hand, binds Eif4e to suppress cap-dependent translation (Bah *et al*, 2015). We co-folded both pairs, Eif4e - Eif4ebp2 (Figure 5C, Supplemental Figure 7A) and Eif4e - Eif4enif1 (Figure 5D, Supplemental Figure 7B), using AlphaFold3. The Eif4e - Eif4ebp2 model is consistent with the PDB structure of human EIF4EBP2 with EIF4E (Fukuyo *et al*, 2011) where Eif4ebp2 contacts the N-terminus of Eif4e. This binding site of Eif4e is the same region where Eif4enif1 is predicted to bind, suggesting mutually exclusive interactions between these two proteins (Figure 5E). We observed a large stem.MAP Confidence Score Delta between the Eif4ebp2-Eif4e interaction and the Eif4enif1-Eif4e interaction (Confidence Score Delta = 0.983), suggesting, in mouse embryonic stem cells, Eif4e is inhibited by Eif4bp2 resulting in cap-independent translation. Further, when examining the profiles of these proteins in our mESC CF-MS data, we observed co-elution between Eif4e and Eif4ebp2. Alternatively, while Eif4enif1 is highly expressed, Eif4e and Eif4enif1 do not co-elute (Figure 5F, Supplemental Figure 7C). This is in contrast with HEK293T CF-MS data, which show Eif4e co-eluting with Eif4enif1 as well as Eif4ebp2 (Figure 5G, Supplemental Figure 7D). Taken together, our observations suggest that Eif4ebp2-Eif4e interactions play roles in mESC cap-independent translation while Eif4enif1 is still expressed and primed to participate in cap-dependent translation upon differentiation.

Oxidative phosphorylation (OXPHOS) has been shown to be highly regulated in embryonic stem cells (Garge *et al*, 2026). We identify two OXPHOS proteins Ndufaf3 and Ndufa6 to be mutually exclusive when binding a common protein Ndufs3. Ndufs3 and Ndufa6 interact in Complex I of the electron transport chain (Guo *et al*, 2017). Ndufs3 also co-immunoprecipitates with Ndufaf3, a Complex I assembly factor, in human cells (Zurita Rendón & Shoubridge, 2012). Further, when Ndufaf3 is genetically disrupted, Ndufs3 is not integrated into Complex I (Murari *et al*, 2021). Structural predictions of Ndufs3 and Ndufaf3 (Supplemental Figure 8A) suggests Ndufaf3 likely binds near the C-terminal region of Ndufs3, similar to the binding mode of Ndufa6 with Ndufs3 (Supplemental Figure 8B). Overlaying these two structural models (Supplemental Figure 8C) highlights the shared binding site of Ndufa6 and Ndufaf3. The models’ predicted aligned error (PAE) plots also indicate that Ndufa6 and Ndufaf3 bind similar regions on Ndufs3 (Supplemental Figure 8D-E). We next asked if mESC prefers one binding partner over another. stem.MAP scores the Ndufs3-Ndufa6 interaction with much higher confidence (stem.MAP score = 0.987) than the Ndufs3-Ndufaf3 interaction (stem.MAP score = 0.01). This is supported by elution profiles of mESC SEC CF-MS data that shows co-elution between Ndufs3 and Ndufa6, but not Ndufaf3 (Supplemental Figure 8F). Further, in differentiated cells such as HEK293T, we observe co-elution of Ndufs3, Ndufa6, and Ndufaf3 (Supplemental Figure 8G) suggesting this effect is specific to mESCs.

### stem.MAP predicts interactions in stem cells independent of priors

To identify potentially novel interactions in mESC, we evaluated the relationship between stem.MAP confidence score and the priors from hu.MAP3.0. Figure 6A shows the majority of confident stem.MAP protein interactions have high hu.MAP3.0 priors suggesting most interactions are not specific to stem cells. Interestingly, we observed 1,388 confident stem.MAP protein interactions without priors (stem.MAP confidence score >= 0.9), suggesting these pairs may be specific to mESCs. We modeled these pairs with AlphaFold3 and identified 28 confident interactions (Figure 6B, Supplemental Table 7, ipSAE >= 0.4). Additionally, we compared our pairs identified with no hu.MAP3.0 priors to an independent network based on quantitative proteomics-based co-regulation derived from naïve and primed ESCs, and early and late gastruloids in human and mouse (Garge *et al*, 2026). We found that 63 of these pairs were highly co-regulated in this independent dataset (Supplemental Table 8). Within this set, we found several uncharacterized interactions. For example, we find Dusp12, a dual specificity protein phosphatase linked to tRNA modifications (Muñoz-Alonso *et al*, 2000), interacts with Kti12, a t-RNA dependent ATPase associated with the Elongator tRNA modification complex (Krutyhołowa *et al*, 2019). Both stem.MAP and AlphaFold3 confidently predict their interaction (stem.MAP score = 0.90, ipSAE = 0.42, Figure 6C, Supplemental Figure 9A). Supporting this prediction, we see consistent co-elution between the proteins in all three orthogonal CF-MS data sets (Figure 6D) and high co-abundance in the gastruloid co-regulation network (Garge *et al*, 2026) (Figure 6E). Dusp12 therefore may regulate several tRNA modification steps (see Discussion).

**Figure 6.**
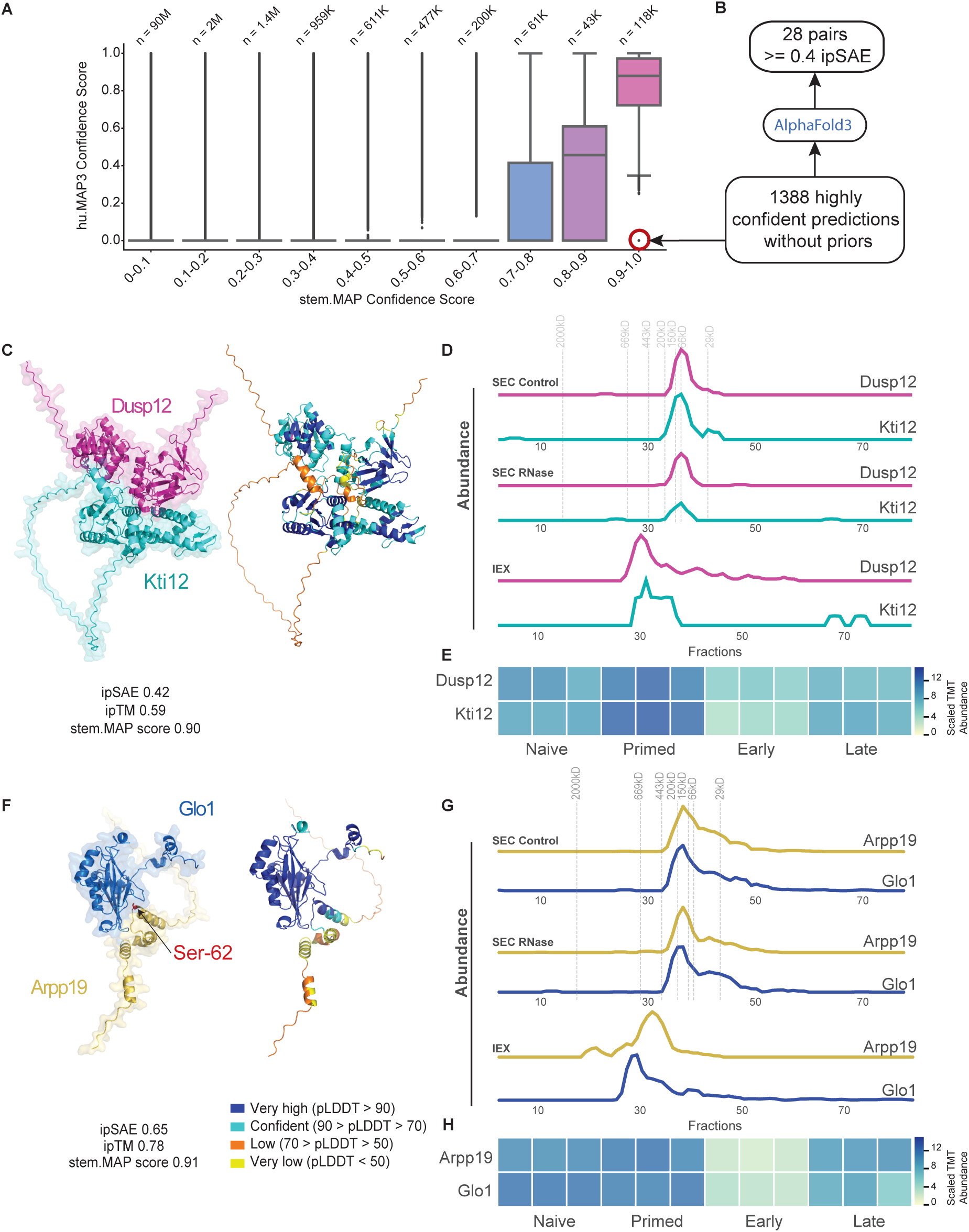
Structural modeling of cell-type-specific stem.MAP predicted PPIs. **A.** Box plots show the relationship of hu.MAP3.0 confidence scores for protein pairs to stem.MAP confidence scores. 1,388 protein interactions predicted by stem.MAP at a confidence of 0.9 and above without hu.MAP3.0 priors are circled in red. The center line of the box plot represents the median, and the upper and lower boundaries of the boxes represent the first and third quartile. **B.** Workflow of selecting high confidence structural models from the protein pairs predicted without hu.MAP 3.0 priors. **C.** AlphaFold3 structures of Dusp12 (pink) and Kti12 (cyan). **D.** Elution profiles of Dusp12 and Kti12. **E.** Protein abundance values of Dusp12 and Kti12 from Garge et al., in naïve mESC, primed mESCS, early and late gastruloids. **F.** AlphaFold3 structures of Arpp19 (yellow) and Glo1 (blue). Serine 62 of Arpp19 is highlighted. **G.** Elution profiles of Arpp19 and Glo1. **H.** Protein abundance values of Arpp19 and Glo1 from Garge et al., in naïve mESC, primed mESCS, early and late gastruloids.

stem.MAP also predicts a confident interaction (stem.MAP score= 0.91) between an inhibitor of the Pp2a complex, Arpp19 and lactoylglutathione lyase, Glo1. AlphaFold3 predicts Arpp19 and Glo1 to have a confident structural interface (ipSAE = 0.65, Figure 6F), and both proteins have consistent co-elution profiles across all three CF-MS experiments (Figure 6G). Further, the proteins show co-abundance in the gastruloid co-regulation network (Figure 6H). When phosphorylated, Arpp19 inhibits the Pp2a complex during the M-phase of mitosis (Gharbi-Ayachi *et al*, 2010). The AlphaFold3 model of Arpp19 and Glo1 shows high pLDDT and confident PAE localized at Ser-62 on Arpp19 (Figure 6F, Supplemental Figure 9B), which is a known phosphorylation site important for entry into mitosis (Gharbi-Ayachi *et al*, 2010). Taken together, this suggests Glo1 may play a role in mitosis by way of its interaction with Arpp19.

stem.MAP also identifies interactions with confident AlphaFold models including a lipase necessary for mESC expansion, Abhd11 (Liu *et al*, 2020), and a carbonic anhydrase, Ca13 (Supplemental Figure 10A). The pair has highly correlated elution profiles (Supplemental Figure 10B) supporting its stem.MAP score of 0.91 and confident PAE plots supporting the AlphaFold model (Supplemental Figure 10C). We also see collagen metabolism related proteins, Serpinh1 (Hsp47), a collagen chaperone important for angiogenesis (Ma *et al*, 2021) and Acp6, a lipid phosphatase whose substrate lysophosphatidic acid, is an important signalling molecule that regulates collagen synthesis (Sato *et al*, 2004). The AlphaFold3 models for this pair also show a high ipSAE and correlated elution profiles (Supplemental Figure 10D-F)

## Discussion

Cellular functions are carried out by proteins that assemble into macromolecular complexes. Distinct cell types in an organism are defined by their various functions and linking these cellular functions to their underlying molecular machinery is a current challenge of molecular biology. We lack easily deployable approaches to identify protein complexes in the diversity of cell types that make up an organism. Here, we describe a high-throughput proteomics workflow based on CF-MS and machine learning to identify cell-type-specific protein complexes (Figure 1) and apply it to mESCs. Our resource, stem.MAP, substantially increases the accuracy and coverage of identified protein complexes in embryonic stem cells. Further, our work represents several advances to gain cell type resolution in protein complex maps (Figure 2). The addition of priors, in particular, allows for an order of magnitude less experiments to be collected, thereby increasing the scalability of our approach.

We show that the resulting protein interaction networks are accurate, identifying many complexes consistent with known biology (Figure 3). We also show our high confident predictions are consistent with protein co-abundance data from Aydin et al. (Aydin *et al*, 2023). We do however see low overall correlation between protein interaction and co-abundance datasets (Pearson correlation coefficient = 0.04) including a mixture of both high co-abundant and high confident stem.MAP pairs as well as low co-abundant and high confident stem.MAP pairs. This result is consistent with previous observations (Fischer *et al*, 2025). It is difficult to rule out technical errors in calculating co-abundance and stem.MAP confidence scores as the reason for the low correlation between the two. Alternatively, a potential biological reason for the low correlation is that accessory and mutually exclusive protein subunits with limited co-abundance to their complexes play an important role in embryonic stem cell function. We identify stem cell associated transcription factors as members of protein complexes (Supplemental Figure 3). In particular, we see Klf4 as a member of a genome integrity complex. Klf4 has been shown to regulate resection and homologous recombination (HR) upon double stranded break repair (Checa-Rodríguez *et al*, 2020) which is the favored pathway by mouse embryonic stem cells over nonhomologous end joining (NHEJ) (Tichy *et al*, 2010). Slf2 is also involved in HR through its recruitment of SMC5/6 to DNA break sites (Räschle *et al*, 2015). Tp53bp1 inhibits HR (Bunting *et al*, 2010), which points to potential crosstalk between the pathways.

We demonstrate our approach is capable of identifying cell-type-specific variants of protein complexes important for stem cell function including the PRC2 complex (Figure 4), BAF, NuRD complexes (Supplemental Figure 4), and Eif4e (Figure 5). Notably, stem.MAP identifies specific components of the embryonic variant of the chromatin remodeling BAF complex (esBAF) including Smarca4(Brg1), Arid1a(BAF250a), and Smarcc1(Baf155). Smarca4 is an ATPase important for removing nucleosomes. It is recruited by Oct4 for chromatin remodeling to allow expression of pluripotency associated genes (King & Klose, 2017), and establishes chromatin for STAT3 signaling (Ho *et al*, 2011). Arid1a, also a member of the esBAF complex, has been shown to contribute to proper expression of ES self-renewal genes including Sox2 and Oct4 (Gao *et al*, 2008). Smarcc1, a scaffolding protein, is critical for esBAF assembly (Schaniel *et al*, 2009) and when depleted in ES cells results in limited proliferation, and apoptosis (Ho *et al*, 2009). These subunits are all highly mutated in cancer cells and neurodevelopmental disorders demonstrating their importance beyond ES cells (Valencia *et al*, 2023). stem.MAP also identifies subunits critical for ES function in the NuRD complex including Mbd3, Cdk2ap1, Hdac1, and Hdac2. Mbd3 is a bridge scaffolding protein found in most cells but when disrupted in ES cells impairs ES differentiation (Kaji *et al*, 2006). Cdk2ap1 is an ES subunit found only in higher order NuRD assemblies (Bode *et al*, 2016) and is necessary for silencing of Oct4 during stem cell differentiation (Deshpande *et al*, 2009). Hdac1 and Hdac2 are deacetylase enzymes which when both are knocked out in ES cells result in loss of cell viability (Jamaladdin *et al*, 2014). The disruption of these enzymes largely affects rapidly dividing cells such as mESC.

Understanding these cell-type-specific subunits is still an active area of research. In our analysis we categorize subunits into “Prototypic”, “ES specific”, and “non-ES specific” classes based on what is found in the functional literature as well as protein expression datasets available for ES cells. Importantly, many of the proteins labeled “Prototypic” are often still critical for ES function but are also important to other cell types (e.g. Mbd3). We expect our resource will aid in linking those subunits important for ES function to their ES specific protein complexes.

Finally, we highlight our workflow’s ability to identify new interactions (Figure 6). For example, stem.MAP identified a confident interaction between Dusp12 and Kti12, supported by AlphaFold models, and co-regulation within a gastruloid network. Kti12 was previously shown to be associated with critical tRNA modifications by the Elongator complex (Scherf *et al*, 2025)) and to bind directly to the elongation factor Elp1 in a phosphorylation dependent manner (Abdel-Fattah *et al*, 2015). Dusp12 also has been associated with tRNA modification by regulating Nat10, a cytosine acetyltransferase which acetylates tRNAs among other targets (Sharma *et al*, 2015; Boell *et al*, 2026). While these two tRNA modifications by Nat10 and Elongator complex are thought to be independent, Dusp12 may play a role in regulation of both with its phosphatase activity.

Similarly, another pair without priors predicted by stem.MAP, and a part of the cell cycle machinery is Arpp19 and Glo1. Arpp19, phosphorylated at a singular Ser-62 residue by Greatwall kinase, is then acted on by the Pp2A complex. The slow dephosphorylation of Arpp19 keeps the Pp2A complex inhibited, and hence Cdk1 and Cyclin-B1 active during the M-phase. A short G1 phase with constitutively active cyclinB1 and Cdk1 is a hallmark cell cycle signature of stem cells (Coronado *et al*, 2013; Kim *et al*, 2018). We saw from our AlphaFold models that Glo1 binds to the Ser-62 residue of Arpp19. This could imply a potential mechanism by which Arpp19 is sequestered from Greatwall Kinase through its interaction with Glo1.

While stem.MAP covers a large portion of the proteome, some limitations of this method still remain. CF-MS experiments are biased towards well-expressed, stable, and soluble protein interactions in cells. For example, Nanog and Myc have low abundance in some CF-MS datasets. While we still identify interactions for these core stem cell factors, we may be missing some interactions due to their low abundance. Overall, we do see excellent recovery of known ES subunits co-eluting with important complexes for stem cell function (e.g. PRC2 and BAF). However, CF-MS is ultimately a bulk measurement of complexes present in a heterogeneous mixture of millions of cells. This may limit the resolution of identifying complexes important for cell-state transitions. Further, the use of priors in our machine learning workflow may bias identification of protein interactions present in other cell types at the expense of ES cell-type-specific protein interactions. We mitigate this by fine-tuning our machine learning pipeline to randomly remove priors from training examples to allow the model to learn examples without such data. As a result, we see high confidence protein interactions predicted by stem.MAP which do not have priors (Figure 6) and are therefore potentially stem cell-specific interactions. Given these limitations, our stem.MAP pipeline accurately identifies protein complexes that are cell-type-specific and can be extended generally to other cell types with limited amount of experimental effort.

## Materials and Methods

### Benchmark curation

Protein complexes for test and training sets were generated as described in (Fischer *et al*, 2025). In brief, human protein complexes were obtained from Complex Portal (version 2022_07_11 https://ftp.ebi.ac.uk/pub/databases/intact/complex/2022-07-11/) (Balu *et al*, 2025). The complexes were split equally into test and training sets. Isoforms and large protein complexes (>30 subunits) were removed. In cases where protein complexes have shared pairs of proteins, one complex was removed from either the test or the training set to prevent data leakage. These sets of protein complexes were then used to generate labeled protein pairs for training and evaluation. Protein pairs from the same protein complex were considered as an interacting pair and labeled as positive. Protein pairs from different protein complexes were considered noninteracting and labeled as negative. The training and test sets had a high negative-to-positive ratio. To limit class imbalance during training, a subset of 10,000 negatives were randomly chosen from all negatives for the training set. Due to our use of hu.MAP3.0 priors, we used the same test and training partition from hu.MAP3.0 to avoid data leakage from the priors during stem.MAP training. The human protein pairs were mapped to their mouse orthologs using inParanoidDB9.0 (Persson & Sonnhammer, 2023) (Supplemental Table 9). To identify proteins of highest similarity, we selected ortholog groups that had a seed score of 1.0 and an inparalog score of 1.0. Proteins with one-to-one mouse orthologs were retained while proteins that showed gene expansion in mice were eliminated. The benchmark resulted in 2,767 train positives, 9,753 train negatives from 378 training complexes and 2,782 test positives, 237,820 test negatives from 378 test complexes (Table 1). All stem.MAP test and training complexes and protein pairs can be found in (Supplemental Table 10) and also on our HuggingFace repository (see Code and Data Availability).

**Table 1:** Number of test and training benchmark interactions.

|  | **Complexes** | **Positive labels** | **Positives labels with feature matrix entries** | **Positives labels with feature matrix entries and >= 0.1 average correlation** | **Negative labels** | **Negative labels with feature matrix entries** |
|---|---|---|---|---|---|---|
| **Test** | 378 | 2,782* | 2,519 | 1,817 | 237,820* | 216,523 |
| **Training** | 378 | 2,767 | 2,487 | 1,452 | 9753 | 8,827 |

\* Evaluation was done on the full test set.

### Dataset collection and preparation

Orthogonal mESC specific CF-MS datasets were collected from Mallam et al. 2019 (SEC) (Mallam *et al*, 2019) and Cox et al. 2026 (IEX) (Cox *et al*, 2026). The detailed methods can be found in the respective papers. The SEC MS data was accessed from ProteomeExchange (PXD014607), processed as described in Mallam et al. Mouse proteome ‘.fasta’ files were downloaded from the UniProt database (2023-05-16; UniProtKB reviewed/SwissProt entries). Raw MS files were converted to mzXML files using MSConvert (http://proteowizard.sourceforge.net/tools.shtml). We used MS-Blender (Kwon *et al*, 2011) to integrate different peptide search engines MSGF+ (Kim & Pevzner, 2014), X! TANDEM (Craig & Beavis, 2004), and Comet (Eng *et al*, 2013). We executed the MS-Blender workflow using a nextflow pipeline using a ‘.config’ file (https://github.com/KDrewLab/stemMAP/tree/main/nextflow) used for peptide identification at a false discovery rate of 1%. Final peptide spectral matches (PSMs) of each protein in each fraction was reported. The.elut files for the IEX data were downloaded from PRIDE (PXD041915) (Supplemental Table 11).

`‘‘‘`

`./nextflow/nextflow run./run_msblender/msblender.nf-with-docker./msblender:latest -- results_path.`

`‘‘‘`

### Feature matrix construction

The collected mass spectrometry data was used to derive features indicative of protein-protein interactions. Only proteins that were determined to have greater than 2 PSMs in at least one of the experiments (i.e., SEC-Control, SEC-RNAse, or IEX experimental datasets) were considered. Pairwise Pearson Correlation scores between two proteins were calculated using the extract_features.py script from the protein complex maps GitHub repository (commit: eebe6c3) (https://github.com/KDrewLab/protein_complex_maps/tree/master/protein_complex_maps/features/ExtractFeatures/canned_scripts) and combined to an initial feature matrix using build_featmap.py script modified from the leca_proteomics repository (commit: f3689d2) (https://github.com/KDrewLab/leca-proteomics/blob/stem_map/update-build-featmat/scripts/build_featmat.py).

`‘‘‘`

`python./extract_features.py MES_SEC_control.elut-f pearsonR-r poisson_noise-i 100-t 2`

`‘‘‘`

This script was run on a directory (e.g../feat_dir/) containing all three extracted Pearson correlation.txt files produced by the extract_features.py script and were given a column name with suffixes “_Mallam_RNAse”, “_Mallam_Cntl”, “_MES_IEX” respectively.

`‘‘‘`

`python build_featmat.py-d./feat_dir/-s _MES_SEC_RNase _MES_SEC_Control _MES_IEX`

`‘‘‘`

To incorporate existing knowledge of protein-protein interactions, we utilized hu.MAP3.0 interaction scores as priors. We first mapped the human protein pairs to their mouse orthologs using inParanoidDB9.0 (Persson & Sonnhammer, 2023) (Supplemental Table 9). To generate prior-weighted correlation scores, we multiplied the hu.MAP3.0 confidence scores by the experimentally-derived individual correlation scores.

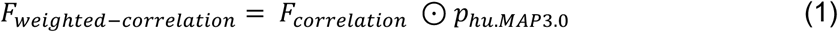

Equation 1 represents the component-wise multiplication of F_correlation_ and p_hu.MAP3.0_, where F_correlation_ is an n x 3 matrix representing correlation coefficients from 3 CF-MS experiments across n protein pairs, and p_hu.MAP3.0_ is an n x 1 vector representing hu.MAP3.0 confidence scores across n protein pairs. F_correlation_ and F_weighted-correlation_ are then concatenated to form an n x 6 matrix representing the full feature matrix. The vector p was padded with random small numbers ranging from 1e-6 to 9e-6 for protein pairs which did not have hu.MAP3.0 confidence scores. The final feature matrix is of the form protein pairs x features, with six total feature columns: three experimentally-derived correlation scores and three prior-weighted correlation scores, and covers 96,618,838 total protein pairs. We provide the full feature matrix on our HuggingFace and Zenodo repositories. We evaluated each of the features using precision-recall curve analysis (https://github.com/KDrewLab/protein_complex_maps/blob/master/protein_complex_maps/evaluation/pl ots/prcurve.py) on the leave-out test set to evaluate their predictive signal (Supplemental Figure 2A).

`‘‘‘`

python./protein_complex_maps/protein_complex_maps/evaluation/plots/prcurve.py -- results_wprob humap3_weighted_pearsonR_MES_SEC_control.pairWpearson humap3_weighted_pearsonR_MES_SEC_RNase.pairWpearson humap3_weighted_pearsonR_MES_IEX.pairWpearson pearsonR_MES_SEC_control.pairWpearson pearsonR_MES_SEC_RNase.pairWpearson pearsonR_MES_IEX.pairWpearson --labels Prior_Mallam_control Prior_Mallam_RNase Prior_IEX_mixedbed Mallam_control Mallam_RNase IEX_mixedbed --input_positives

`./Mouse_mapped_IntAct_TRAIN_featmat_humap2_bioplex3_WMM_trimCols_groupbyMean_from20 220625_20230503_FILTERED_positive_interactions_20231026.txt --input_negatives`

`./Mouse_mapped_IntAct_TRAIN_featmat_humap2_bioplex3_WMM_trimCols_groupbyMean_from20 220625_20230503_FILTERED_negative_interactions_20231026.txt --output_file`

`./PR_Curve_test_MES_individual_features.pdf --complete_benchmark --add_tiny_noise-`

`-precision_markers 0.5 0.1`

`‘‘‘`

### Pre-training engineering

To make the benchmark more cell-type-specific, we removed positive pairs from the training set with low correlation scores in the mESC CF-MS data. First, for each positive pair, we averaged the Pearson correlation values across all three CF-MS experiments. We then determined the percent of positive pairs of the training set which had lower average correlation values for a range of cutoffs (0, 0.1, 0.2, 0.3, 0.4, 0.5) (Supplemental Figure 1). Based on this analysis we chose to remove positive training pairs with average correlation values < 0.1, as this preserved 1,452 of training positives with data (Supplemental Figure 1). The negative training pairs were unaltered.

### Building the stem.MAP model

To construct a model to identify protein-protein interactions, we utilized AutoGluon (v0.8.2) (Erickson *et al*, 2020), an open-source machine learning framework to train a weighted-ensemble classifier. Autogluon efficiently explores a range of independent random forest, gradient boosting, and combinations of model ensembling to optimize predictive performance. During training, we utilized balanced accuracy as our evaluation metric and trained the following base models: XGBoost, LightGBM, RandomForestGini, NeuralNetTorch, NeuralNetFastAI, ExtraTreesGini, ExtraTreesEntr, RandomForestEntr, LightGBMLarge, CatBoost, KNeighborsUnif, LightGBMXT, KNeighborsDist. All models were evaluated for their performance on a 10% hold-out validation set.

Only ∼20% of protein pairs with data in our feature matrix also have priors from hu.MAP3.0. To build a model which can accurately predict interactions for the 80% of pairs without priors, we randomly replaced priors from the training feature matrix with a small random number between 1e-6 to 9e-6. To select the percentage of training priors to replace for final model training, we built models with 0%, 5%, 10%, 20%, 30%, and 40% of random training priors replaced. We then evaluated the trained models with the full test set and a test set with only 20% of the protein pairs with priors representing the real world scenario. The 10% model was chosen because it had consistent performance with the model with no priors replaced on the full test set and superior performance on the test set representing the real world scenario (Figure 2F). We evaluated the stem.MAP model using precision-recall analysis on the leave out test set (https://github.com/KDrewLab/protein_complex_maps/blob/master/protein_complex_maps/evaluation/pl ots/prcurve.py). The top performing model was a weighted ensemble model, consisting of LightGBM and NeuralNetTorch with weights 0.667 and 0.333 respectively and was chosen as our final stem.MAP model. Additionally, we evaluate model confidence using a precision vs confidence score analysis (Supplemental Figure 2B) and compare with hu.MAP3.0.

### Clustering and parameter selection

We used a two-stage clustering approach as described in (Fischer *et al*, 2025) to identify protein complex clusters within the protein interaction network. We thresholded the network based on stem.MAP confidence score of 0.1 and above. To identify the dense regions of the network we used ClusterOne (Nepusz *et al*, 2012) and to each dense region we applied the MCL (Enright *et al*, 2002) algorithm to fine tune the clustering. We used the network_clustering.nf nextflow script from the protein complex maps repository for generating clusters of different MCL and ClusterOne parameters (Commit eebe6c3) as specified by the.config file: https://github.com/KDrewLab/protein_complex_maps/blob/stem_map/protein_complex_maps/clustering/network_clustering.config. We performed parameter sweeps across model confidence score thresholds (0.987, 0.986, 0.985, 0.984, 0.983, 0.982, 0.981, 0.98, 0.96, 0.95, 0.93, 0.91, 0.9, 0.8, 0.7, 0.6, 0.5, 0.4, 0.3, 0.2, 0.1), ClusterOne density (0.3, 0.35, 0.4), ClusterOne max overlap (0.6, 0.7, 0.8), and MCL inflation (1.2, 2, 3, 4, 5, 7, 9, 11, 15), totaling 1,701 parameter combinations.

`‘‘‘`

./nextflow run./network_clustering.nf

`‘‘‘`

Cluster sets were evaluated using weighted average precision and weighted average recall (Drew *et al*, 2017, 2021) against the training set of gold standard protein complexes from complex portal (See Benchmark curation in Methods) using the psweep_comparison.py script (commit 3045ac1) https://github.com/KDrewLab/protein_complex_maps/blob/stem_map/protein_complex_maps/evaluatio n/psweep_comparison.py.

`‘‘‘`

python./psweep_comparison.py --cluster_filenames./*.nr.txt --gold_standard

./mouse_mapped_complex_portal.train.txt --output_filename

./stemMAP_clustering.psweep_comparison.txt --proc 8 --normalize_by_combinations

`‘‘‘`

We chose two clusterings, one exhibiting high precision and one exhibiting high recall and made a final complex map from the union of both the clusters (Supplemental Table 12). No duplicate complexes were found between the high precision and high recall cluster map. Protein complexes with over 100 subunits were removed from the final map. Final evaluation against the Complex Portal leave-out test set yielded a weighted average precision of 0.44 and weighted average recall of 0.39 (Supplemental Figure 2E).

### Ablation tests

Ablation tests were done on the final stem.MAP weighted ensemble model with a modified test feature matrix. Two test feature matrices were built with either the Pearson correlation scores, or the prior-weighted correlation scores replaced with zeros. A third feature matrix was built where all the features were retained but the values within a column were shuffled, to assess how the model performed on randomized data. The predictions of the pre-trained stem.MAP model on the modified test feature matrices were evaluated on the test set using a precision-recall curve.

### Annotation enrichment of stem.MAP protein complexes

We used g:Profiler(Kolberg *et al*, 2023) to calculate Gene Ontology enrichment for all the complexes in stem.MAP. All proteins with at least 2 PSMs in any one of the experiments (n=14,896) were considered as background proteins. We calculated enrichments from GO, HP, CORUM, REAC, and KEGG. We randomly shuffled the complexes while preserving the cluster size of each complex and compared the shuffled enrichments against stem.MAP complex enrichments under a p-value of 0.01 (Supplemental Table 4).

### Embryonic stem cell-specific annotations and associated proteins

Embryonic stem (ES) cell associated (ES-associated) proteins were collected from (Ng & Lufkin, 2011) (https://pmc.ncbi.nlm.nih.gov/articles/PMC3360086/table/T2/), Oct-4 and Nanog pull down experiments (van den Berg *et al*, 2010; Wang *et al*, 2006; Pardo *et al*, 2010), and mESC-unique proteins from (Van Hoof *et al*, 2006). These are highlighted in the cytoscape network (Zenodo) and in Figure 3D. These proteins were used to annotate proteins in stem.MAP to identify ES-associated interactions and complexes. For mESC-unique and mDiff-unique proteins (Supplemental Figure 3D) we collected proteomics data for mESCs and their differentiated derivatives from (Van Hoof *et al*, 2006). We removed any overlapping proteins expressed between mESCs and the differentiated derivatives. We also compared stem.MAP to a naïve mESCs proteomics dataset (Atlasi *et al*, 2020), which is closer in developmental state to the mESCs used for CF-MS from Mallam et al and Cox et al. We obtained the standardized counts for protein abundances of naïve mESCs from PAX-DB (Wang *et al*, 2012) for the (Atlasi *et al*, 2020) dataset. (https://pax-db.org/downloads/6.1/datasets/10090/10090-PXD014528_proteome_esc.txt). We retained only proteins that were more than 10 NSAFs (normalized spectral abundance factor). Overlapping proteins between Atlasi et al mESC proteins and Van-hoof et al mESC-unique proteins were taken together as mESC consensus proteins and overlapping proteins from Atlasi et al in Van Hoof et al in the differentiated-derivative proteins were removed. The remaining set was taken as mDiff consensus proteins. The average abundance of mESC consensus and mDiff consensus proteins were calculated from the three CF-MS datasets and were compared (Supplemental Figure 3E).

To classify subunits of complexes as ES, non-ES, and Prototypic subunits we utilized a combination of proteomics literature as well as literature based on functional experiments to determine relevance to stem cell self renewal and pluripotency. ES subunits were defined as those interacting in the complex exclusively in stem cells or whose loss impairs pluripotency, self-renewal, or causes lethality in stem cells. To define subunits of the Baf complex we used (Zhang *et al*, 2021; Gatchalian *et al*, 2018; Ho *et al*, 2009; Alfert *et al*, 2019; Gao *et al*, 2008; Zhang *et al*, 2019). For the NuRD complex, we utilized (Mor *et al*, 2018; Bode *et al*, 2016; Schmolka *et al*, 2023; Zhao *et al*, 2017; Deshpande *et al*, 2009; Liao *et al*, 2023). For the PRC2 complex, we utilized (Piunti & Shilatifard, 2021; Chamberlain *et al*, 2008; Kloet *et al*, 2016). Mbd3, Chd4, and Eed are shown to have important roles in embryonic stem cells, they have roles in other cell types and therefore were marked as prototypic subunits.

### Coabundance of proteomic measurements from outbred mouse embryonic cell line

Normalized protein abundance values from 190 genetically diverse mouse embryonic cell lines was downloaded from here: https://pmc.ncbi.nlm.nih.gov/articles/instance/10112288/bin/mmc2.xlsx (Aydin *et al*, 2023). Pearson correlation coefficients were calculated using the normalized protein abundance values for all protein pairs identified in the study. A two tailed KS test (mpmath stats (Johansson *et al*, 2017)) was performed to compare the distributions of co-abundance between protein pairs with stem.MAP confidence scores >= 0.8 and < 0.8.

### Mutually exclusive protein pairs analysis

A list of 5,781 mutually exclusive interactions predicted between a protein trimer was obtained from the hu.MAP3.0 publication (Fischer *et al*, 2025) (https://static-content.springer.com/esm/art%3A10.1038%2Fs44320-025-00121-5/MediaObjects/44320_2025_121_MOESM8_ESM.xlsx). The pairs were mapped to mouse and 3,789 groups with both mouse orthologs and stem.MAP score were used for our analysis. We calculated the absolute difference between the stem.MAP scores and the hu.MAP3.0 scores of each pair of mutually exclusive interactions using the following equation:

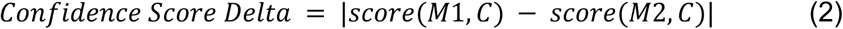

where M1 is mutually exclusive protein 1 interacting with the common subunit C and M2 is mutually exclusive protein 2 interacting with the common subunit C.

### AlphaFold3 structural modeling

While hu.MAP3.0 scores provided priors for many of the protein pairs in this network, stem.MAP also identified 1,388 pairs of proteins without prior hu.MAP3.0 scores as likely to interact. To add more orthogonal evidence to this set, we ran systematic pairwise model prediction using AlphaFold3. Pairwise models were generated using the official public release of AlphaFold3 (Abramson *et al*, 2024) (github.com/google-deepmind/alphafold3) installed locally, as well as the public AlphaFold3 server.

Input consisted of only the two peptide chains in a given stem.MAP pair, with no post-translational modifications or ligands. AlphaFold3 server runs were performed without manual seeding, equivalent to selecting a random integer as a seed. A model seed of 1 for all predictions performed locally. Computation was performed for local AlphaFold3 runs on Indiana University’s Jetstream2 HPC cluster. To make effective use of resources across CPU and GPU cluster environments, the multiple sequence alignment and model inference stages of the AlphaFold3 pipeline were separated utilizing its -- norun_inference and --norun_data_pipeline flags. Aside from these, AlphaFold3 predictions were run with default settings. 1342/1388 pairs produced AF3 models. The remaining models failed due to insufficient memory. AF3 structures of human mutually exclusive pairs are from (Jänes *et al*, 2024; Burke *et al*, 2023), AF3 structures of the mouse mutually exclusive proteins represented in Figure 5 and Supplemental Figure 8 were modeled from the AF3 web server. The models were evaluated using the interaction prediction Score from Aligned Errors (ipSAE) (Dunbrack, 2025) using the ipsae.py script from the IPSAE repository on GitHub (https://github.com/DunbrackLab/IPSAE). A PAE cutoff of 10Å and distance cut off of 10Å was used for ipSAE calculation. Using performance evaluation done with 40 known binding proteins with PDB structures calculated with these parameters in (Dunbrack, 2025), we chose a ipSAE confidence cutoff of >= 0.4. All ipSAE calculated scores for structural models are reported in Supplemental Table 7.

## Code and Data Availability

Training and test feature matrix, stem.MAP confidence pairwise confidence scores can be found in HuggingFace repository (model: https://doi.org/10.57967/hf/11343, data: https://doi.org/10.57967/hf/11342). All code can be found on github repository (https://github.com/KDrewLab/stemMAP). stem.MAP complexes, network, feature matrix, and benchmarks can be found on the downloads page of our web resource (https://stem.proteincomplexes.org/download). Annotated cytoscape network, full stem.MAP network, feature matrix, AlphaFold3 models and evaluation metrics, and Pearson correlation coefficients for protein pairs in Aydin et al. can be found on Zenodo (https://doi.org/10.5281/zenodo.22978322).

## Acknowledgements

K.D. and S.F. are supported by the National Institute of Child Health and Human Development (R00 HD092613) and BBSRC – National Science Foundation/Directorate for Biological Sciences [BB/X002179/1][BB/X002683/1][NSF2314278]. K.D., S.F., and M.W.G. are supported by NIGMS (R35GM164016). Research reported in this publication was supported by the National Institute of General Medical Sciences of the National Institutes of Health under Award Number R35GM164016. The content is solely the responsibility of the authors and does not necessarily represent the official views of the National Institutes of Health. Computational resources were provided by the NSF ACCESS Exchange [BIO250091], and the University of Indiana JetStream2. R.K.G. was supported by the Washington Research Foundation, National Human Genome Research Institute (NHGRI; 1RM1HG010461), Eunice Kennedy Shriver National Institute of Child Health and Human Development (NICHD; 1K99HD120632). A.R. was supported by the American Heart Association Pre-doctoral fellowship (24PRE1189649).

## Supplemental Tables

**Supplemental Table 1: Autogluon machine learning model leaderboard**

**Supplemental Table 2: List of stem.MAP complexes**

**Supplemental Table 3: stem.MAP pairwise predictions**

**Supplemental Table 4: Complex annotation enrichment**

**Supplemental Table 5: Aydin et al co-abundance scores**

**Supplemental Table 6: List of preferred mutually exclusive partners**

**Supplemental Table 7: AF3 structural model confidence scores**

**Supplemental Table 8: Gastruloid co-regulation network correlation score for pairs without priors**

**Supplemental Table 9: Human to mouse Inparanoid mapping**

**Supplemental Table 10: Test and Training benchmark sets**

**Supplemental Table 11: mESC CF-MS elution files**

**Supplemental Table 12: Clustering parameters**

## Supplemental Figure Legends

**Supplemental Figure 1.**
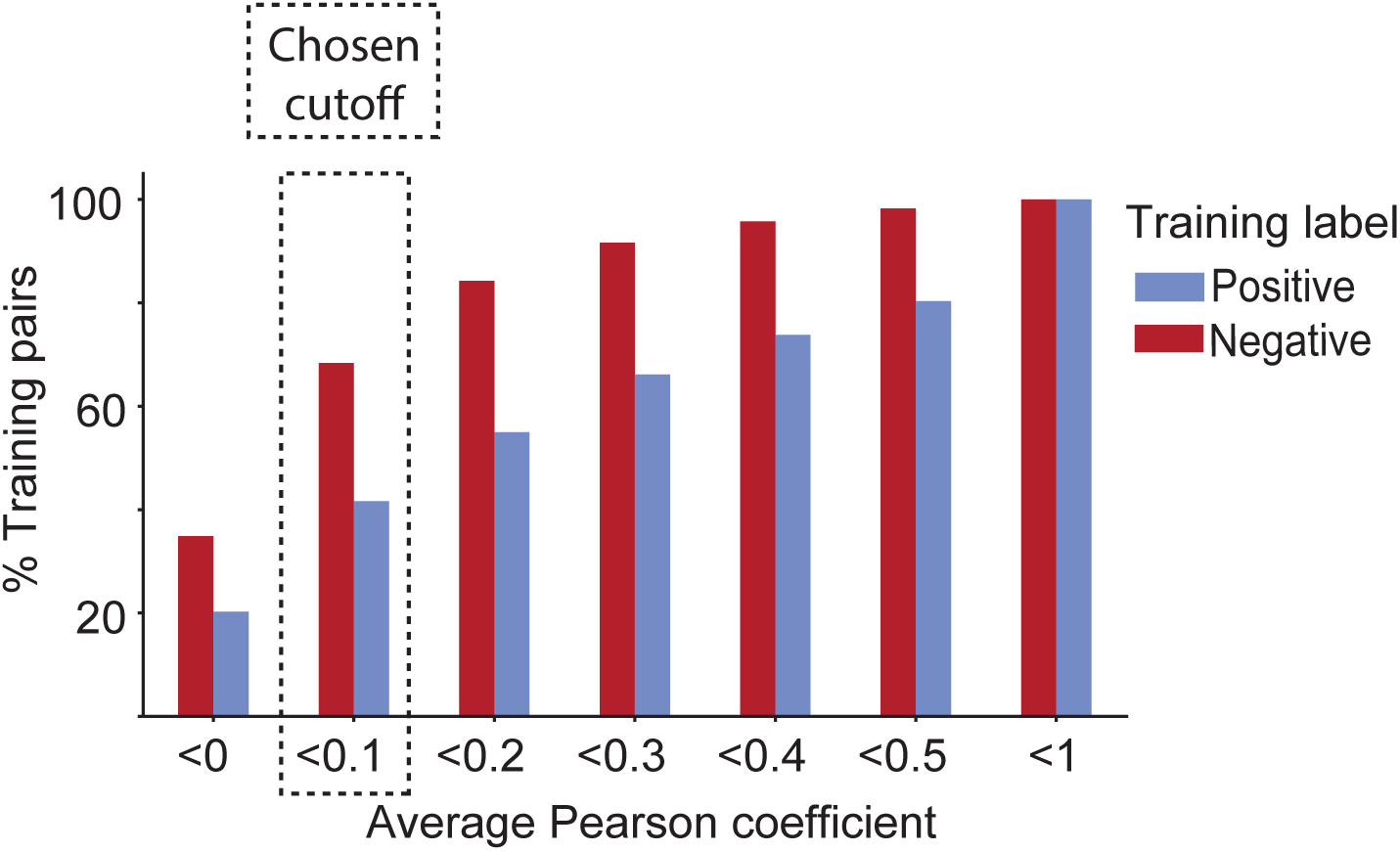
Selection of pre-training engineering benchmark cutoff. Percent of positive training labels with data and average Pearson correlation coefficients less than specified cutoff. Percent of negative labels are shown for reference.

**Supplemental Figure 2.**
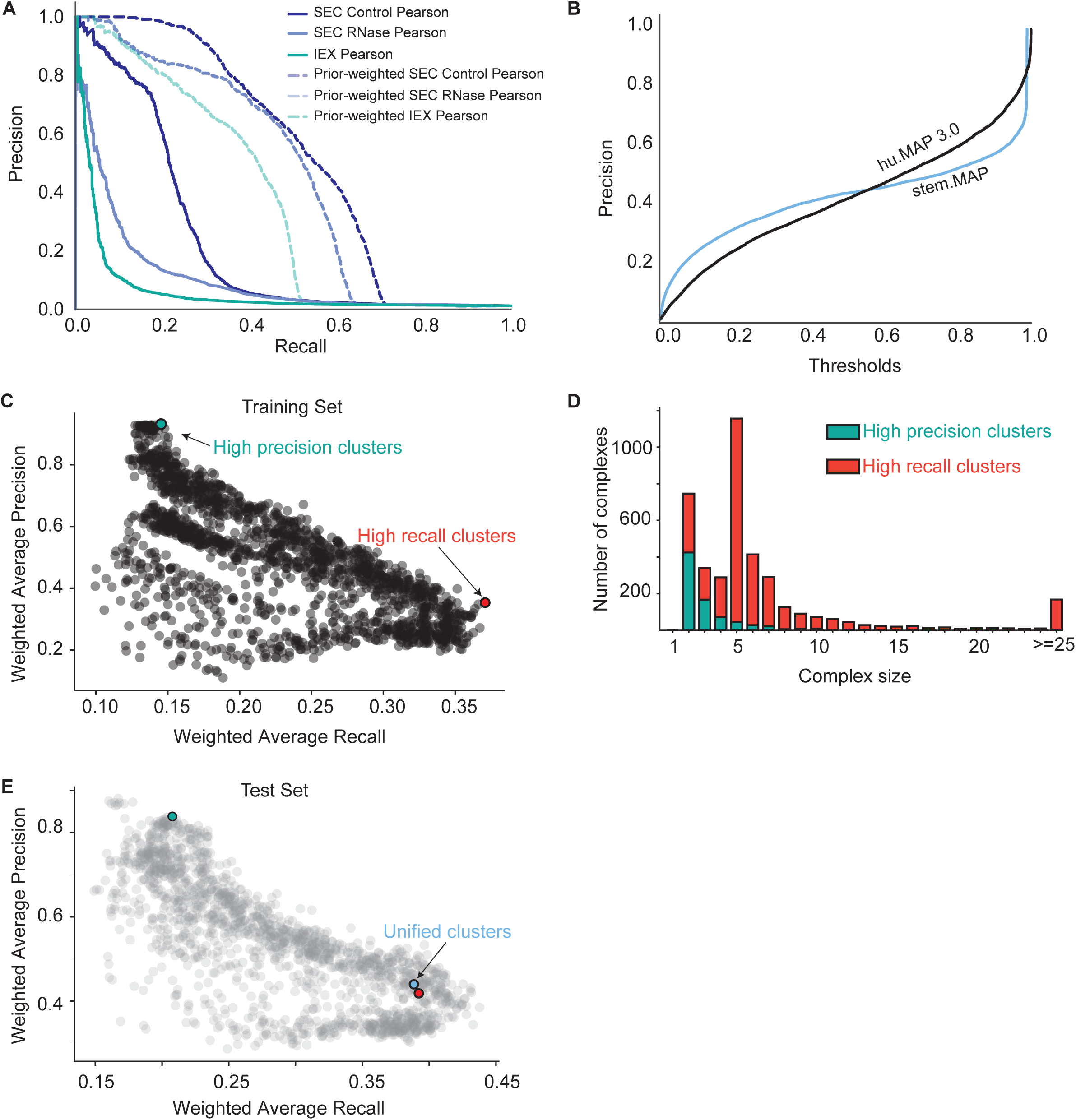
stem.MAP evaluation and clustering optimization. **A**. Precision-vs-recall curves evaluating pairwise Pearson correlation coefficient (solid lines) from each CF-MS experiment (Mallam et al SEC control, SEC RNase treated and Cox et al., IEX respectively) with their corresponding prior weighted correlation (dashed lines). **B**. Precision-vs-threshold curve evaluating stem.MAP and hu.MAP3.0. stem.MAP confidence score of 0.8 is at a precision of approximately 0.5. **C**. Clustering performed using different MCL and ClusterONE parameters were evaluated using weighted average precision vs weighted average recall scatter plot on the training set of complexes as detailed in hu.MAP cluster evaluation. A high precision (teal) cluster and a high recall cluster (red) were chosen as final sets of clusters. **D**. Stacked histogram comparing complex size vs number of complexes for both the high precision and the high recall cluster group. **E**. High precision (teal), high recall (red) and union (blue) clusters were compared to the rest of the clustering parameters (gray) using weighted average precision and recall on the test complexes.

**Supplemental Figure 3.**
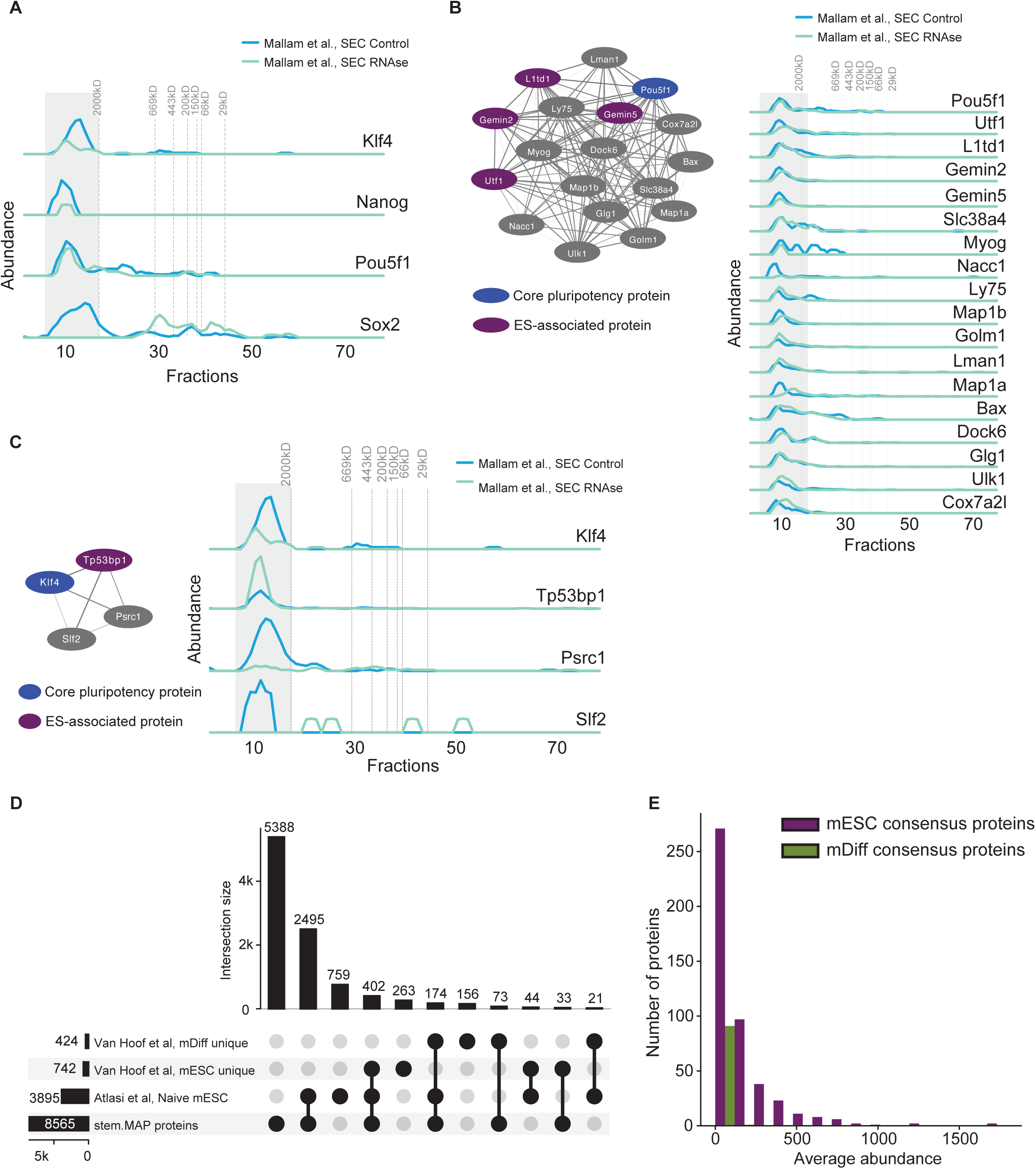
mESC new interactions predicted with the core pluripotency transcription factors. **A**. Sparklines from SEC control and SEC RNase datasets for Klf4, Nanog, Pou5f1 (OCT4 in humans), Myc, and Sox2. **B**. Complex containing Pou5f1 pluripotency factor (blue) (stemMAP_00115.1). ES-associated proteins (purple) predicted in the complex include L1td1, Gemin2, Gemin5 and Utf1. Elution profiles in SEC CF-MS experiments show high co-elution of subunits. Edge weights are proportional to stem.MAP confidence score. **C**. Complex containing Klf4 pluripotency factor (blue) (stemMAP_01109.1). Elution profiles of SEC CF-MS experiments show high co-elution of subunits. **D**. UpSet plot showing the overlap of proteins in stem.MAP with mESC unique mDiff unique proteins described by Van Hoof et al., naïve mESC proteins described in Atlasi et al. **E**. Average abundance of mESC (purple) and mDiff (green) consensus proteins (see Methods), identified in the mESC CF-MS datasets used in stem.MAP.

**Supplemental Figure 4.**
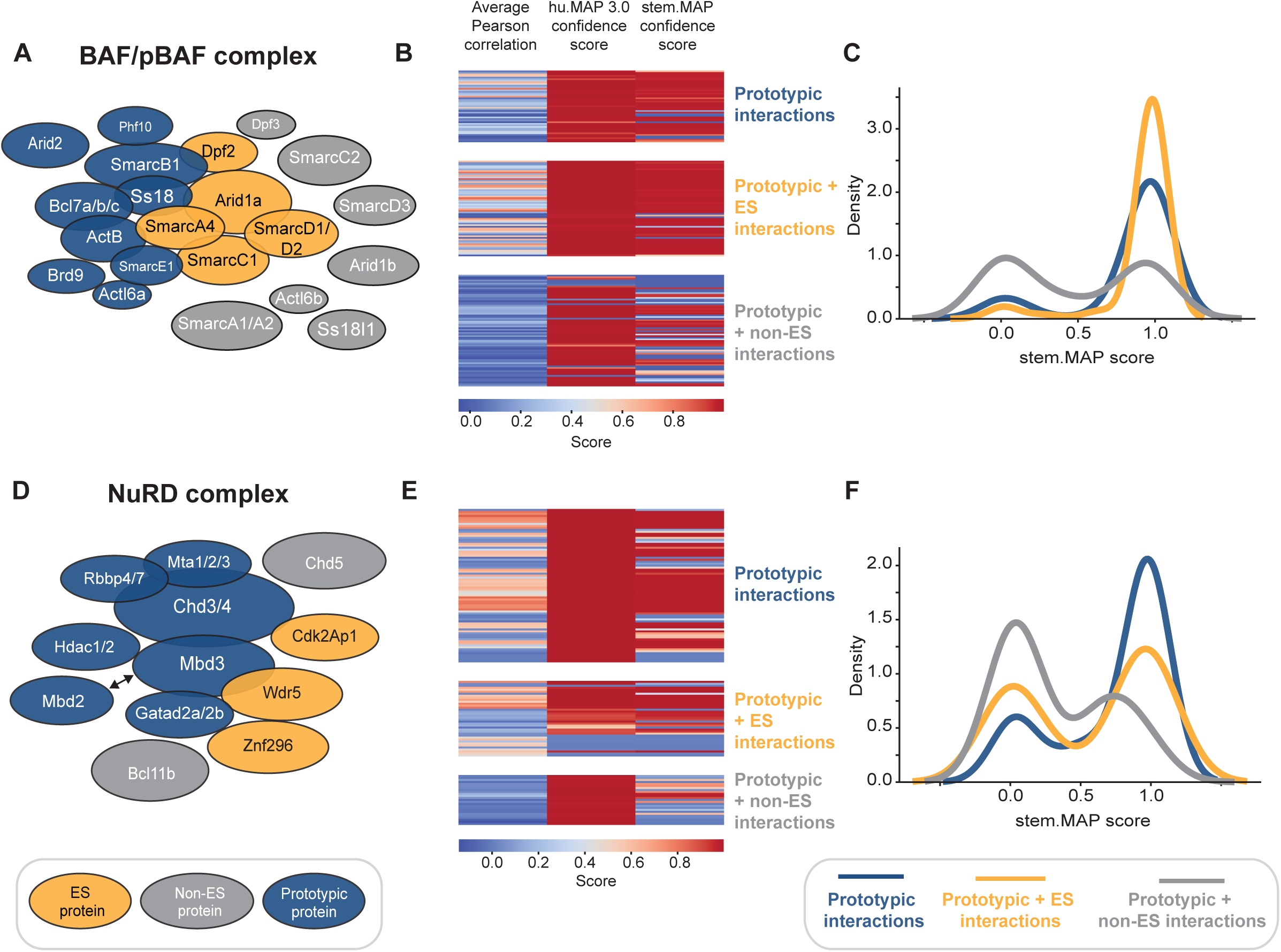
mESC specific interactions in protein complexes predicted by stem.MAP. **A**. Literature annotated BAF complex. Prototypic subunits (dark blue) are defined as proteins that are found in different cell types including embryonic stem cells. ES subunits (yellow) are proteins seen primarily in mESCs as a part of a complex and/or essential for ES self renewal and pluripotency. Non-ES subunits (gray) are proteins found in other cell types. **B** Pairwise scores between prototypic subunits, prototypic + ES subunits, and prototypic + non-ES subunits. Heatmaps represent the average pairwise Pearson correlation score, the pairwise hu.MAP3.0 confidence score used as a prior, and the final pairwise stem.MAP confidence score. **C**. Distributions of stem.MAP confidence scores for the three categories, prototypic pairs, prototypic + ES pairs, and prototypic + non ES pairs. stem.MAP produces high confidence scores for prototypic (blue) and prototypic + ES (yellow) pairs (large right-shifted density), and low confidence scores for prototypic + non-ES specific (gray) pairs (left-shifted density). **D**. Literature annotated NuRD complex. Colors are the same as **A**. **E**. Pairwise scores for NuRD subunit pairs. Descriptions are the same as **B**. **F**. Distributions of stem.MAP confidence scores for NuRD complex. Descriptions are the same as **C**.

**Supplemental Figure 5.**
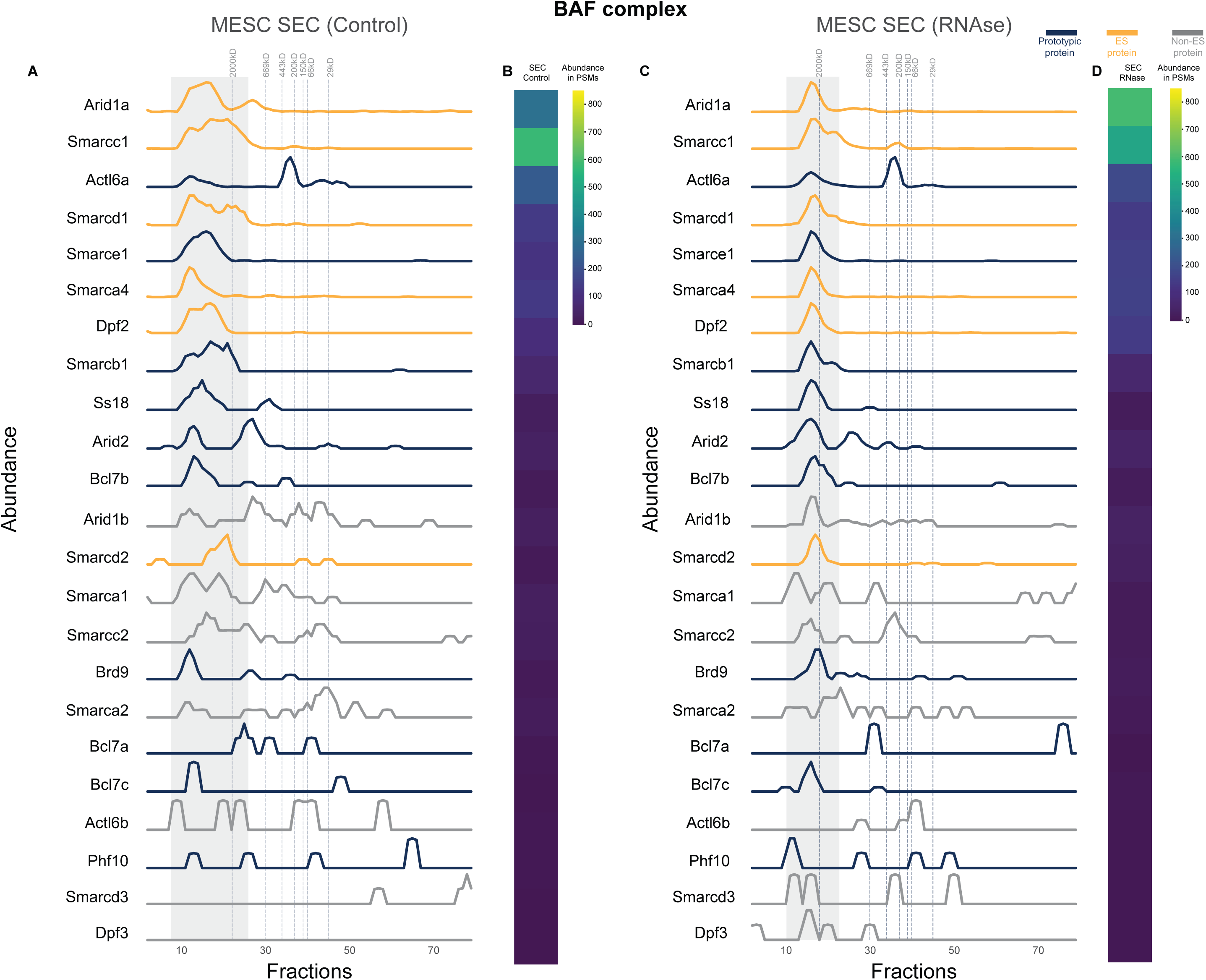
Elution profiles for the BAF complex. **A**. Elution profiles of BAF complex proteins from mESC SEC CF-MS control experiment. **B**. PSM (peptide spectral match) abundance values for each subunit as the sum of all fractions in mESC SEC CF-MS control experiment. **C**. Elution profiles of BAF associated proteins from mESC SEC SF-MS RNase-treated experiment. **D**. PSM (peptide spectral match) abundance values for each subunit as the sum of all fractions in mESC SEC CF-MS RNase-treated experiment. mESC-specific proteins and proteins involved in pluripotency or self-renewal (yellow), proteins specific to a differentiated cell type (gray), and proteins expressed in multiple cell types including mESC (blue). Subunits Ss18l1 and Actb were not detected in the SEC experiments. Proteins are sorted based on their average abundance in the 3 CF-MS experiments from highest to lowest.

**Supplemental Figure 6.**
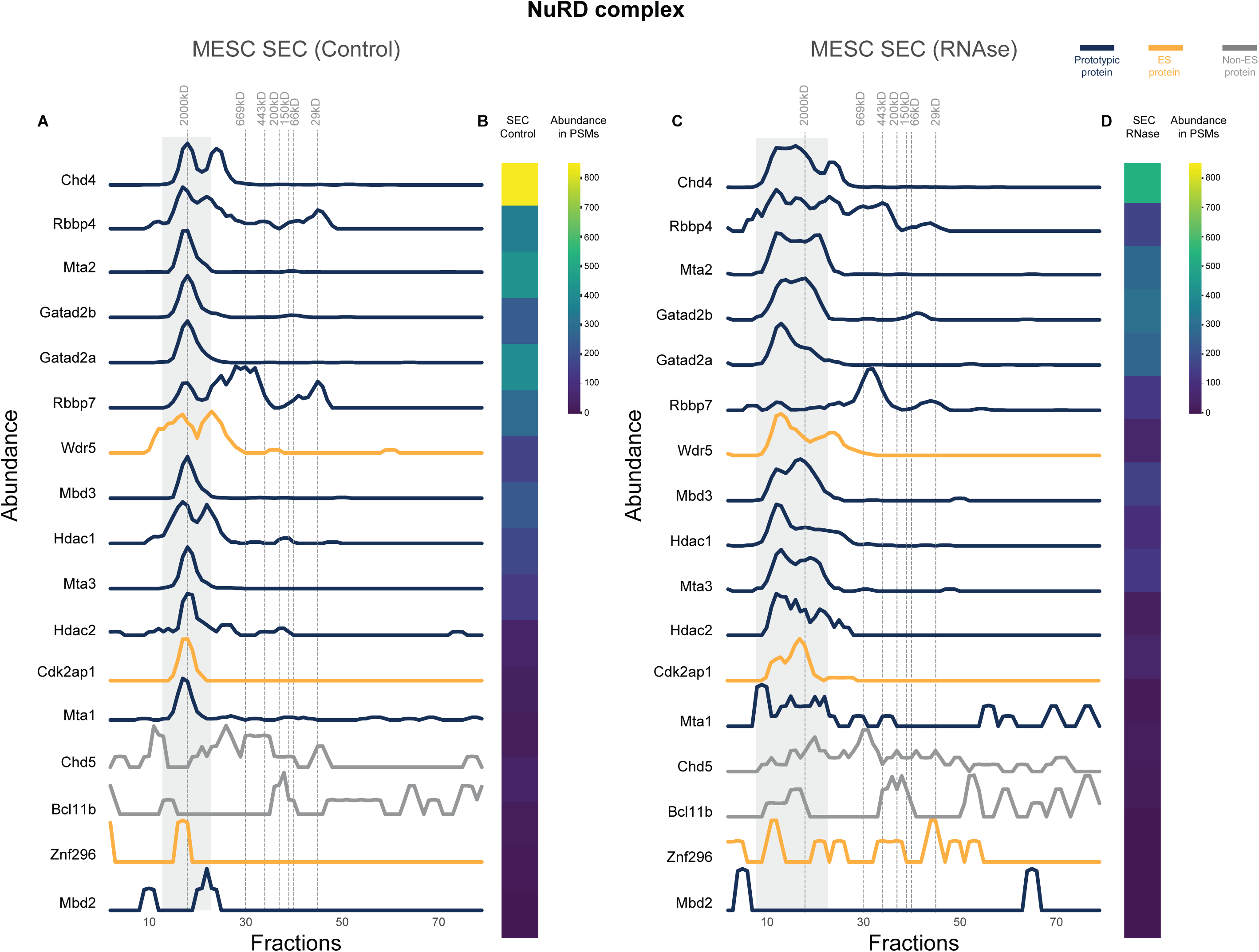
Elution profiles for the NuRD complex. **A**. Elution profiles of NuRD associated proteins from mESC SEC CF-MS control experiment. **B**. PSM (peptide spectral match) abundance values for each subunit as the sum of all fractions in mESC SEC CF-MS control experiment. **C**. Elution profiles of NuRD associated proteins from mESC SEC SF-MS RNase-treated experiment. **D**. PSM (peptide spectral match) abundance values for each subunit as the sum of all fractions in mESC SEC CF-MS RNase-treated experiment. mESC-specific proteins and proteins involved in pluripotency or self-renewal (yellow), proteins specific to a differentiated cell type (gray), and proteins expressed in multiple cell types including mESC (blue). Prototypic subunit Chd3 was not detected in the SEC experiments. Proteins are sorted based on their average abundance in the 3 CF-MS experiments from highest to lowest.

**Supplemental Figure 7.**
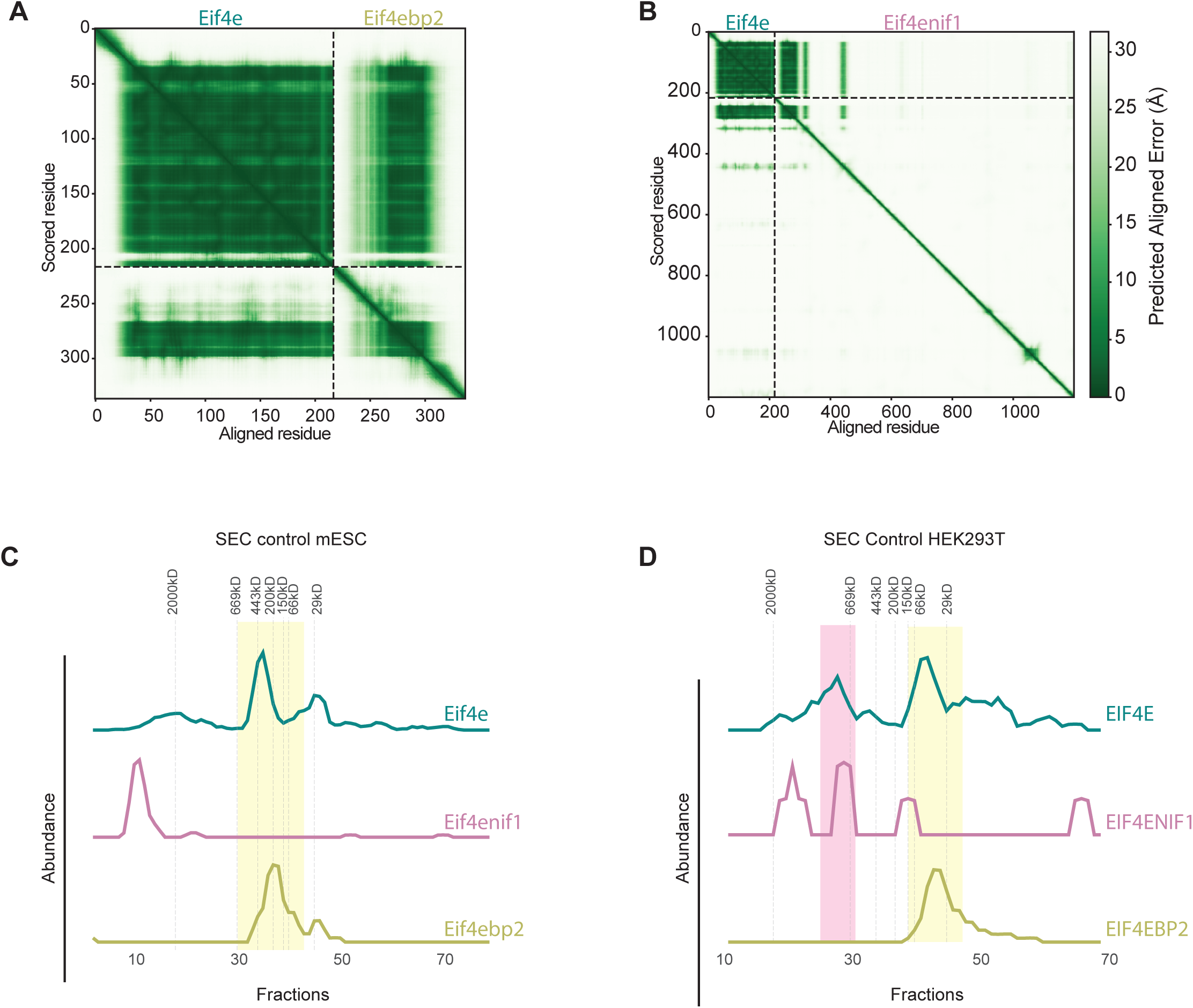
Mutually exclusive interactions with the eukaryotic initiation factor Eif4e - elution profiles and PAE plots: **A**. and **B.** PAE plots of Eif4e with Eif4ebp2 and Eif4enif1 with an ipSAE of 0.83 and 0.829 respectively. **C.** CF-MS elution profiles from mESC SEC control experiments. **D**. CF-MS elution profiles from HEK293T cells. The putative inhibited Eif4e complex with Eif4ebp2 is found in both mESCs and HEK293T cells (yellow highlight), whereas the Eif4e complex with Eif4enif1 imported into the nucleus is found only in HEK293T cells (pink highlight).

**Supplemental Figure 8.**
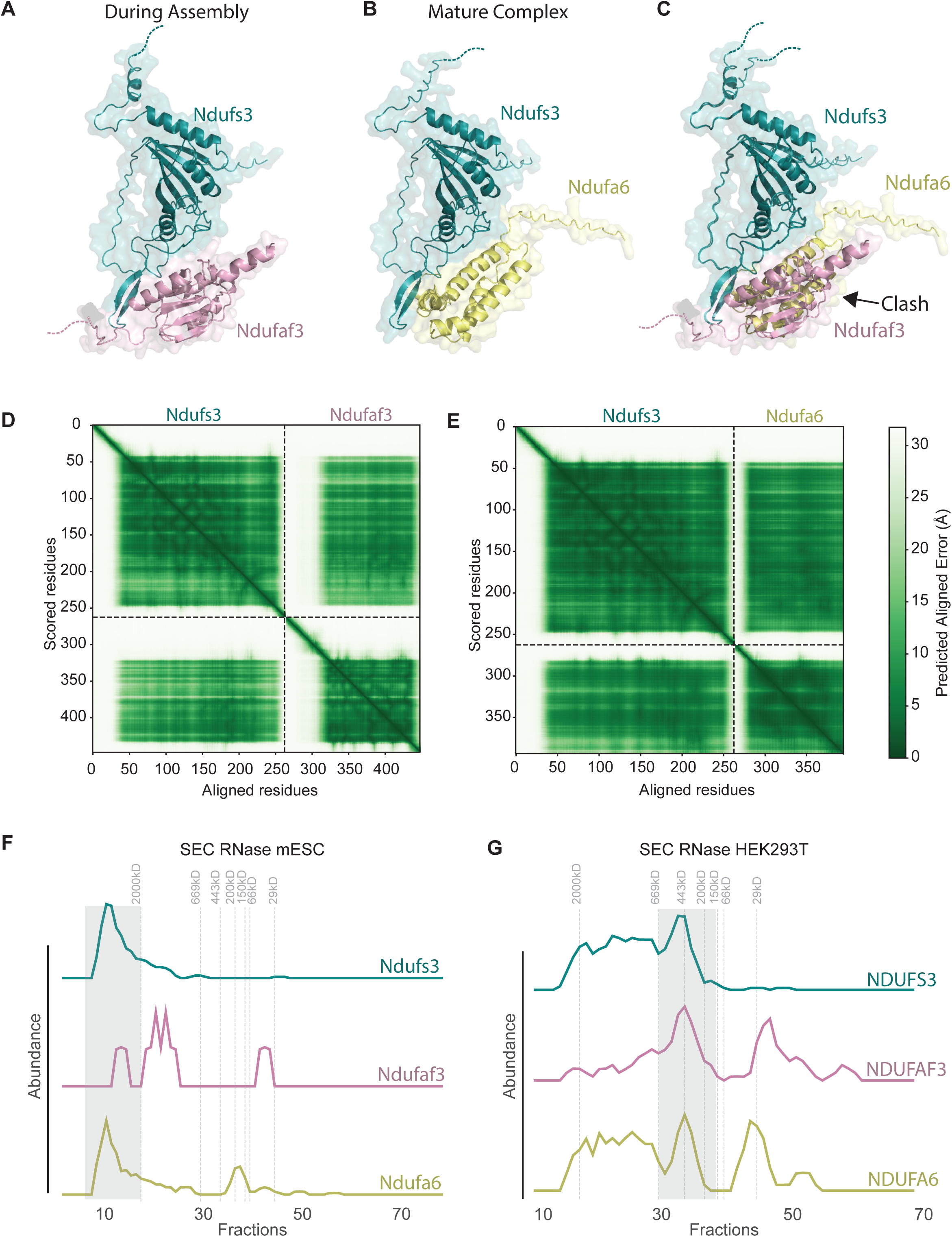
Mitochondrial complex I mutually exclusive interactions of Ndufs3 with Ndufaf3 assembly factor and Ndufa6 structural factor. **A**. AlphaFold model of mitochondrial complex I protein Ndufs3 interacting with Ndufaf3, the chaperone protein for complex assembly. **B**. AlphaFold model of Ndufs3 and Ndufa6, an accessory non-catalytic protein. **C**. Both binding partners of Ndufs3, Ndufaf3 and Ndufa6, occupy the same space causing a clash making them a mutually exclusive interaction pair. **D**. PAE plot of the structural model of Ndufs3 with Ndufaf3, ipSAE = 0.44. **E**. PAE plot of structural model of Ndufs3 and Ndufa6, ipSAE = 0.61. **F**. Elution profiles of Ndufs3, Ndufaf3, and Ndufa6 in mESC SEC CF-MS experiment indicate co-elution of Ndufs3 and Ndufa6 but poor co-elution with the assembly factor Ndufaf3. Gray highlight represents the fully assembled Complex I. **G.** Elution profiles of Ndufs3, Ndufaf3, and Ndufa6 in HEK293T SEC CF-MS experiment indicate co-elution of all three proteins. Gray highlight represents the fully assembled Complex I complex.

**Supplemental Figure 9.**
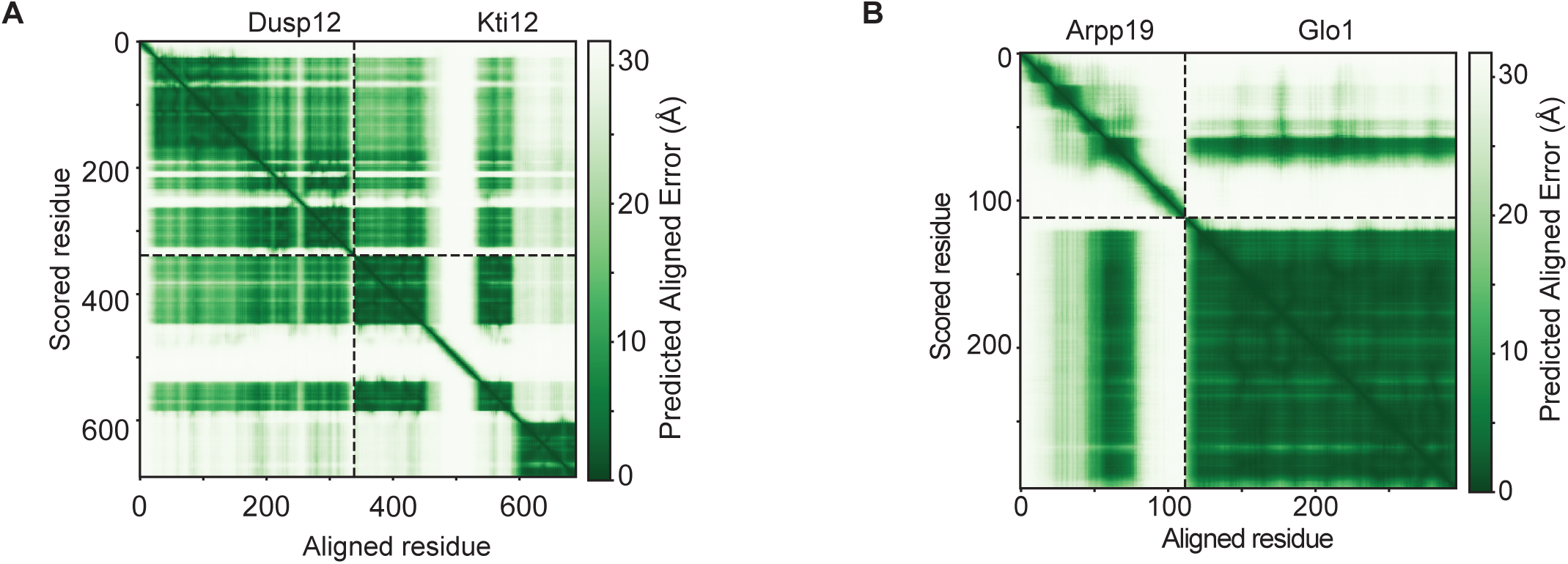
PAE plots of Co-regulation supported AF3 structures. Predicted Aligned Error (PAE) plots of pairwise protein interactions identified from stem.MAP, **A**. Dusp12 and Kti12 and **B.** Arpp19 and Glo1.

**Supplemental Figure 10.**
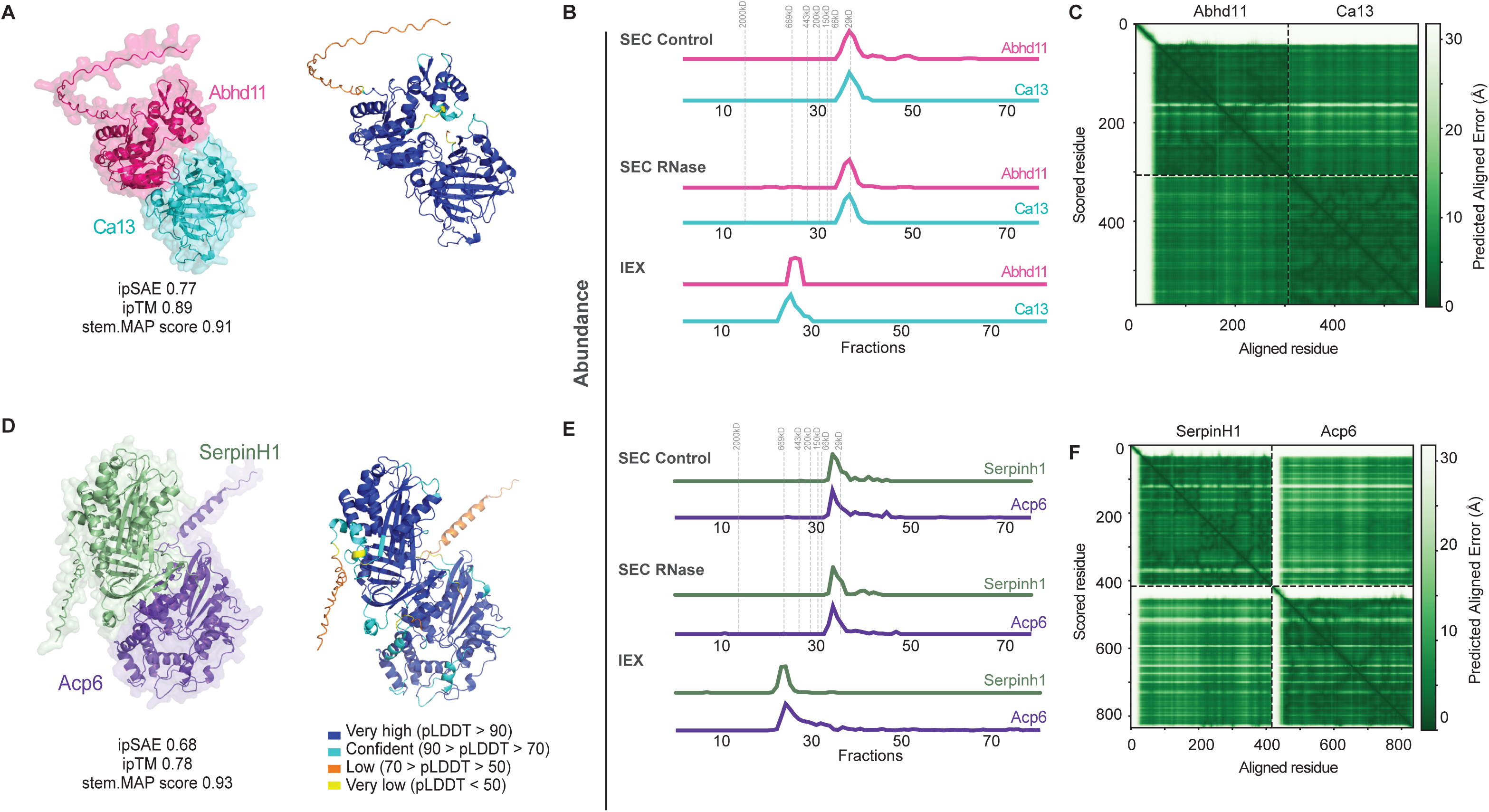
AlphaFold modeling of stem.MAP predicted pairs without hu.MAP3.0 priors A., B., and. **C.** Abhd11 and Ca13. **C.,D., and E.** SerpinH1 and Acp6. (**A. and D.)** AlphaFold3 models are colored by subunits (left) and pLDDT (right). (**B. and E.)** Elution profiles of the protein pairs in SEC-control, SEC-RNase and IEX experiments. (**C and F)** Predicted Aligned Error (PAE) plots of the respective AlphaFold 3 models.

